# Oocyte ETC function and glycogen metabolism links maternal environment to offspring metabolism

**DOI:** 10.1101/2025.05.13.653493

**Authors:** Jason F. Cooper, Umanshi Rautela, Kim Nguyen, Darrick Gates, Emily Wolfrum, Colt Capan, Hyoungjoo Lee, Devia Williams, Chidozie Okoye, Kelsie Nauta, Megan Senchuk, Arlene Garcia, Jian Wei Tay, Ximena Sanchez-Avila, Ryan T. Kelly, Maulik Patel, Ryan Sheldon, Andrew P. Wojtovich, Nicholas O. Burton

**Author notes:** These authors contributed equally to this work (Jason F. Cooper, Umanshi Rautela).

## Abstract

Maternal exposure to environmental stressors can drive adaptive changes in offspring metabolism across diverse species and is linked to disease risk in humans. Such effects have most often been attributed to epigenetic mechanisms, but whether a mother’s environment can also act on other mechanisms present in oocytes to alter offspring physiology remains unclear. Using a system in which *C. elegans* exposure to osmotic stress can drive a greater than 10-fold increase in offspring survival in response to future osmotic stress, we performed global proteomics profiling of dissected oocytes and discovered that maternal exposure to osmotic stress reduces the abundance of electron transport chain (ETC) proteins in oocytes. We further found that ETC function in oocytes promotes oocyte glycogen abundance. Metabolism of this oocyte-derived glycogen, in both oocytes and early embryos, antagonizes the phosphorylation of the AMP-kinase catalytic subunit AAK-2 at Threonine-243 in embryos. Lastly, we find that changes in AAK-2 activity in turn regulate the offspring’s response to future osmotic stress, likely by converging on regulation of the ribosome with the integrated stress response (ISR) pathway. Collectively, our findings describe an intergenerational signaling cascade in which multiple maternal inputs, including maternal environments and maternal insulin signaling, converge on oocyte glycogen metabolism to trigger adaptive changes in offspring physiology via altering AMP-kinase activity in embryos. Notably, recent studies have similarly discovered the existence of insulin-regulated ETC and glycogen remodeling in fly, frog, and human oocytes. However, the function of this evolutionarily conserved phenomenon remains unclear. Our findings suggest that oocyte ETC/glycogen remodeling is regulated by a mother’s environment and, at least in some contexts, links a mother’s environment to adaptive changes in offspring physiology.

## Introduction

For most of the 20th century, organismal responses to environmental conditions were believed to be confined to the individual, with each generation thought to “reset” without inherited information about prior conditions. This belief was largely derived from August Weismann’s influential barrier hypothesis in 1892, which proposed that information about the environment could not be transmitted from somatic cells to germ cells^1^. In the ensuing 100 years, however, parental exposures to common environmental stresses (nutrient availability, infection, osmotic stress) have been reported to drive germ-cell transmissible, stress-specific, and adaptive changes in offspring across diverse evolutionary taxa, including humans^2–9^, where parental and early-life exposures to stress are increasingly linked to offspring growth rates and long-term health outcomes^10–13^. The earliest example to break the Weismann’s barrier hypothesis came from silk moths (*Bombyx mori*), in which maternal external temperature was found to regulate the release of diapause hormone that signals directly to oocytes to trigger changes in offspring sugar metabolism and entry into an overwintering diapause state^14^. The precise mechanisms by which changes in oocytes alter *B. mori* offspring metabolism and diapause entry remain unclear. However in the century since, the mechanisms most often implicated in transmitting a mother’s environment to her offspring via oocytes have been epigenetic, in the form of chromatin marks, DNA methylation and small RNAs-passed through the germline^15,16^.

Studies of germ-cell transmissible, multigenerational effects of stress have often focused on male germ cells (sperm) because this cell type is easy to collect and such effects can be more clearly differentiated from *in utero* effects. By contrast, most mechanistic work on maternal effects have focused on how the fetus or embryo itself is altered by a mother’s environment during gestation, rather than the oocyte, before fertilization. Comparatively little is known about whether a mother’s environment or maternal signaling pathways can directly act on oocytes to alter an offspring’s phenotype. This is in part due to the substantial technical challenges of working with oocytes, which are a relatively rare cell type that largely cannot be cultured *in vitro*. Furthermore, many of the most robust examples of maternally-regulated adaptive changes in offspring metabolism occur in animals that are not easily studied in the laboratory or in animals with relatively long generation times that render mechanistic intergenerational studies impractical^3^ (*e.g.* maternal regulation of offspring growth rates in response to nutrient availability changes in red squirrels^6^). The field also largely lacks unbiased approaches to identify potentially novel mechanisms for information transfer via oocytes, and as a result, most studies have tested *a priori* candidates such as epigenetic marks but have limited ability to identify unexpected mechanisms of information transfer that might uniquely exist in oocytes.

We previously found that *C. elegans*, whose short generation time and genetic tractability make it uniquely suited to dissecting such mechanisms, undergoes a robust oocyte-transmitted adaptation to osmotic stress via a mechanism that requires substantial shifts in offspring metabolism^3^. Interestingly, like the original findings for diapause hormone signaling to oocytes regulating offspring entry into a diapause state in *B. mori*^14^, we found that insulin-like signaling to oocytes regulated the decision of *C. elegans* to enter a diapause like state in response to osmotic stress in *C. elegans*^3^. However, the specific molecular mechanisms present in oocytes that promote this change in offspring remain unknown. Here, we combined *C. elegans* genetics with low-input oocyte proteomics to find that maternal osmotic stress exposure alters the abundance of electron transport chain (ETC) proteins in oocytes and that ETC function in the maternal germline is required for *C. elegans* intergenerational adaptation to osmotic stress. We further found that ETC function in oocytes promotes glycogen accumulation in oocytes and that glycogen abundance in oocytes (and its metabolism in both oocytes and early embryos) is a node that links multiple maternal inputs (maternal environment, maternal insulin signaling) to future offspring stress responses. Lastly, we identify AMP-kinase (AAK-2) as a downstream effector of this pathway in offspring. Maternal disruption of oocyte glycogen metabolism alters phosphorylation of AAK-2 at its canonical regulatory site, Thr243, in embryos and this modification is required for the offspring to adapt to future osmotic stress. Together, these findings establish an intergenerational signaling pathway linking maternal somatic inputs (environment, insulin signaling) to metabolic remodeling of the oocyte, through to a defined molecular response in the offspring.

## Results

### Maternal exposure to osmotic stress alters ETC protein abundance in oocytes

Proteomic studies of fly and frog oocytes have suggested that maternal insulin signaling regulates a previously unappreciated mitochondrial proteome remodeling event in oocytes that includes the reduced abundance of several ETC complexes and results in the functional loss of Complex I^17^. A similar process was also recently observed in human oocytes^18^. Despite these observations across distantly related species, the biological significance of oocyte ETC remodeling remains unclear. Because *C. elegans* intergenerational adaptation to osmotic stress is also regulated by maternal insulin signaling to oocytes^3^, we hypothesized that ETC remodeling might be functionally involved in the process that links maternal exposure to osmotic stress to offspring’s response to future stress.

To test this hypothesis, we first developed and validated a low-input proteomics approach, modified from recently described single-cell proteomics techniques^19^, to reliably quantify the relative abundance of ETC proteins in as few as nine dissected *C. elegans* oocytes per biological replicate. To validate that we can detect changes in the abundance of ETC subunits across conditions using low-input oocyte samples, we first compared the oocyte proteome of wild-type animals to *nduf-7(et19)* mutants, a Complex I subunit whose *et19* mutation is presumed to reduce, but not eliminate, NDUF-7 protein^20^. **(Fig. 1A, Supplemental Fig. 1A, B).** This approach detected peptides matching 8,821 proteins, of which 6,621 passed a ≥ 2 peptide detection filter and 127 exhibited altered abundance in *nduf-7* mutant oocytes compared to wild-type oocytes (**Fig. 1B and Supplemental Table 1**). As expected, NDUF-7 itself was reduced ∼5-fold in *nduf-7* mutant oocytes, accompanied by reduced abundance of multiple other Complex I subunits consistent with Complex I instability^21,22^ (**Fig. 1B**). By contrast, we did not observe significant changes in the abundance of subunits of Complexes II, III, IV or V, suggesting that the changes caused by a mutation in *nduf-7* are specific to Complex I as expected (**Supplemental Table 1**). Notably, cellular component enrichment analysis of the 46 proteins that were reduced in abundance in *nduf-7* mutant oocytes identified Complex I subunits and mitochondrial proteins as the only significantly enriched categories (**Fig. 1C**). We therefore conclude that our low input proteomics approach reliably and specifically detects expected changes in the relative ETC protein abundance from small amounts of oocyte material.

**Figure 1.**
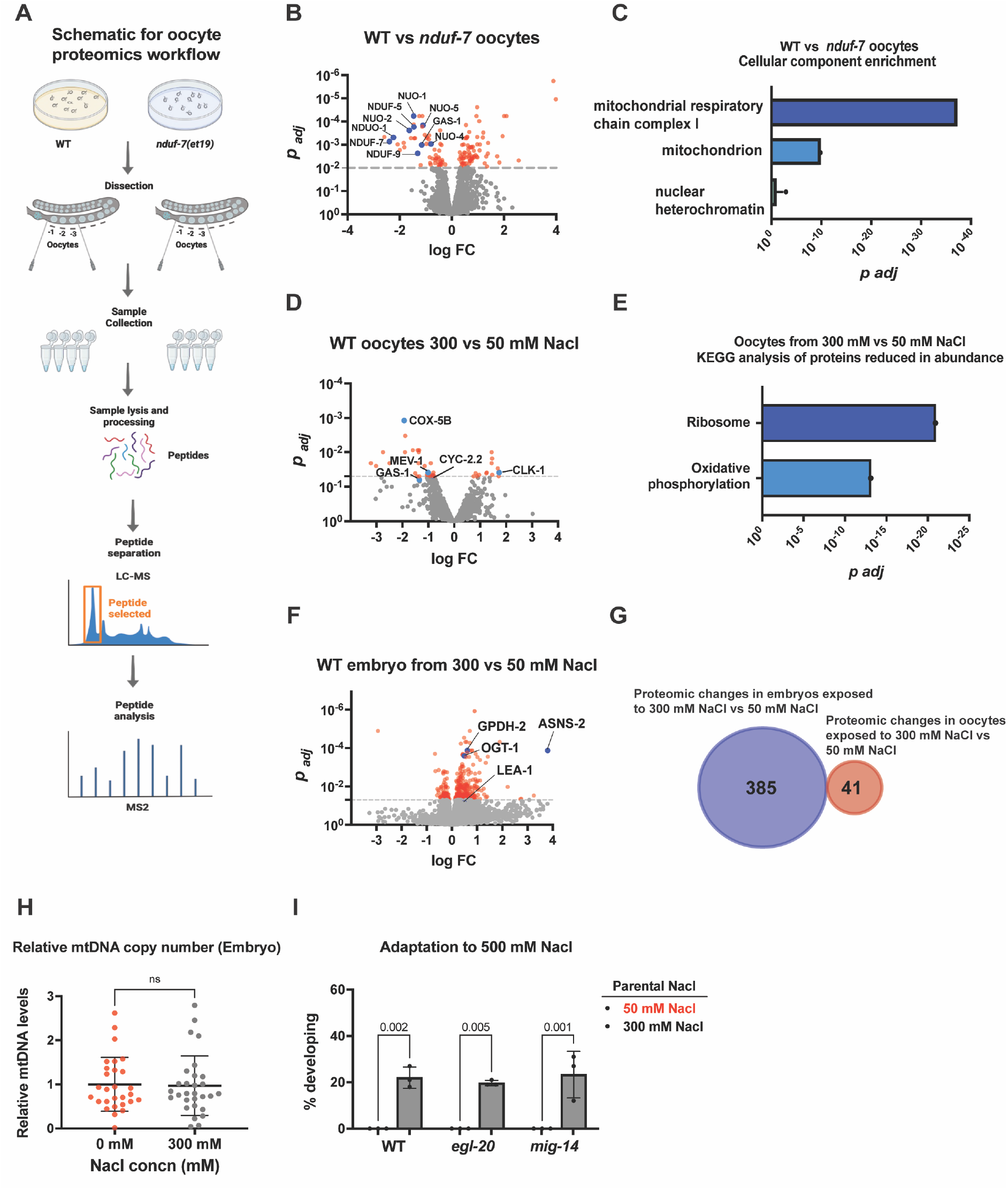
Maternal osmotic stress induces remodelling of ETC protein abundance in oocytes. **A.** Low input oocyte proteomic workflow. Briefly late-L4 staged *C. elegans* wild-type (WT) mothers were grown on control or 300mM NaCl cultures for 24 hours. Seven animals were then isolated, the mature -1, -2, and -3 oocytes were then dissected out of the germline and collected for lysis and sample processing in preparation for low input proteomics via LC-MS. **B.** Volcano plot of 3,123 detected proteins that passed all filters in 3 biological replicates of day-1 wild-type (WT) or *nduf-7*(*et19*) mutant oocytes. Red dots represent proteins that exhibited *p_adj_* < 0.05. Blue dots represent subunits of Complex I. **C.** Biological compartment analysis of proteins altered abundance in *nduf-7*(*et19*) mutant oocytes compared to wild-type oocytes using WormEnrichr ^76,77^. **D.** Volcano plot of 2,866 detected proteins from 6 biological replicates of wild-type oocytes exposed to either 300 mM or 50 mM NaCl for 24 hours. Wild-type control samples at 50 mM NaCl are the same as in Fig. 1B. Red dots represent proteins that exhibited *p_adj_* < 0.05 changes in abundance. Blue dots represent subunits of different complex in ETC. **E.** Biological compartment analysis of proteins altered in abundance in oocytes from parents exposed to 300 mM NaCl compared to wild-type oocytes from parents exposed to 50 mM NaCl using WormEnrichr^76,77^. **F.** Volcano plot of 7,149 proteins in embryos from WT parents exposed to either 50 mM (standard) or 300 mM NaCl (stressed) for 24 hours. Red dots are the 385 proteins that exhibit significant (*p_adj_* < 0.05) changes in abundance relative to the 50 mM NaCl condition. **G.** Venn diagram of proteins that changed in abundance in WT embryos (from animals exposed to 50 mM or 300 mM NaCl for 24 hours) and proteins that changed in abundance in from WT oocytes (from mothers exposed to 50 mM or 300 mM NaCl for 24 hours). **H.** Relative mitochondrial DNA copy number was determined using digital-droplet PCR (ddPCR) in embryos from *C. elegans* mothers exposed to 50 mM NaCl or to 300 mM NaCl (mild osmotic stress) for 24 hours. n - 28 replicates. Unpaired students’ *t*-test with Welch correction. error bars -s.d. **I.** Early day-1 adult staged WT, *egl-20(n585)*, and *mig-14(ga62)* mutant animals were exposed to 50 mM or 300 mM NaCl for 24 hours. Following this the F1 embryos were isolated and exposed to 500 mM NaCl. The number of embryos that developed to larval stage after 48 hours was scored. n = 3 replicates of > 40 offspring per replicate, Two-way ANOVA with Šídák’s multiple comparisons test, error bars – s.d. Source data are provided in Supplemental table 8.

Using this validated method, we next asked whether maternal exposure mild osmotic stress alters the oocyte proteome. Wild-type animals were exposed to either standard laboratory conditions (50 mM NaCl) or mild osmotic stress (300 mM NaCl) after which oocytes were dissected and subjected to global proteomics profiling (**Fig. 1D**). In this comparison, we quantified the relative abundance of 2,866 proteins (**Supplemental Table 3**) of which 29 exhibited significantly decreased abundance and 13 exhibited increased abundance in oocytes from animals exposed to 300 mM NaCl when compared to control oocytes (**Fig 1D, Supplemental Table 2**). Proteins that decreased in abundance were significantly enriched for ribosomal proteins and proteins involved in oxidative phosphorylation (*e.g.* ETC Complex subunits), while those that increased were enriched for mRNA processing proteins, although much less significantly than the change observed for ribosomal and ETC related proteins (**Fig 1E**, **Supplemental Fig 1C**).

Notably, the protein exhibiting the most significant change in abundance was COX-5B, a subunit of Complex IV of the ETC. Similarly, we also observed reduced abundance of GAS-1 (Complex I), MEV-1 (Complex II), F45H10.2 and T02H6.11 (Complex III), and CYC-2.2 (cytochrome c) (**Fig 1D, Supplemental Table 2**). These changes are unlikely to be explained by simple changes in mitochondrial abundance as we did not observe any change in subunits of Complex V (ATP synthase) or in mtDNA copy number in oocytes (**Supplemental Table 3, Supplemental Fig 1D**). Furthermore, the reduced abundance of these ETC subunits persisted even when we normalized abundance to the mitochondrial outer membrane transporter complex, a common proxy for mitochondrial abundance (**Supplemental Table 3**). Together this data suggests a selective reduction in ETC protein abundance rather than a general loss of mitochondrial content. RNA-seq of dissected oocytes further showed that none of the protein-level changes were accompanied by corresponding changes in mRNA abundance (**Supplemental Fig. 1E and Supplemental Table 4**), consistent with the long-standing view that mature oocytes are largely transcriptionally quiescent^23^. We conclude that maternal exposure to osmotic stress selectively reduces the abundance of several ETC proteins in oocytes through a mechanism that is unlikely to involve transcriptional regulation.

Having established that maternal exposure to osmotic stress selectively reduces ETC subunit abundance in oocytes, we next asked whether the reduced ETC protein abundance observed in oocytes was maintained in embryos following fertilization. This is important because persistent changes in the embryo could provide a direct explanation for how maternal stress alters offspring physiology, whereas changes that are restricted to the oocyte would suggest that the oocyte state is subsequently interpreted by the embryo. We therefore performed global proteomic profiling of F1 embryos from parents exposed to either 50 mM NaCl or 300 mM NaCl for 24 hours. We found that 385 proteins exhibited significantly altered abundance in embryos from parents exposed to 300 mM NaCl. These proteins included increased abundance of several proteins known to participate in the osmotic stress response of *C. elegans* (the glycerol-3-phosphate dehydrogenase GPDH-2, the O-GlcNAc transferase OGT-1^23^, the late embryo abundant protein LEA-1^24^) and ASNS-2, a well-established hallmark of the integrated stress response that responds to many different stressors^25^ **(Fig. 1F and Supplemental Table 5**). Despite the increase in abundance of proteins known to be involved in the osmotic stress response, none of the 42 proteins that changed in abundance in oocytes remained altered in abundance in embryos (**Fig. 1G, Supplemental Tables 3 and 5**). Thus, the mitochondrial protein abundance changes observed in oocytes in response to osmotic stress did not persist in the embryonic proteome.

This conclusion also extends to mitochondrial abundance more broadly. Consistent with our results of unchanged mtDNA copy number in oocytes in response to osmotic stress (**Supplemental Fig 1D**), we observed no increase in mtDNA copy number in offspring of stressed parents (**Fig 1H**). We note that a recent study by Zhang *et al.* (2021) showed that increased mtDNA copy number in germ cells could transgenerationally alter *C. elegans* stress responses^26^. As part of these studies, they found that changes in WNT signaling regulated mtDNA copy number and that changes in WNT signaling mediated multigenerational changes in mitochondrial function. However, we found that the same WNT signaling mutants that were found to regulate mtDNA abundance transgenerationally (*egl-20*, *mig-4*) did not affect *C. elegans* adaptation to osmotic stress in our system (**Fig 1I**). These results indicate that *C. elegans* intergenerational response to osmotic stress likely does not act through general changes in mitochondrial abundance (at either the mtDNA or protein level) in oocytes or offspring. Instead, our data point to changes in oocyte ETC function as potentially an independent mechanism that links maternal osmotic stress exposure to offspring’s future stress response.

### ETC function in oocytes impacts offspring’s future response to osmotic stress

While our results suggested that ETC protein abundance in oocytes was dependent on a mother’s environment, whether this ETC remodeling had any functional consequence for the offspring was unknown. Because *C. elegans* intergenerational adaptation to osmotic stress is transmitted via oocytes, we utilized existing genetic mutants with partial loss-of-function mutations in subunits of ETC complexes -*nduf-7(et19)*^20^ and *nuo-6(qm200)*^27^ (Complex I), *mev-1(kn1)*^28^ (Complex II), and *isp-1(qm150)*^29^ (Complex III) – with an aim of testing whether reducing ETC function in oocytes impacts offspring’s future response to osmotic stress. We note here that most ETC null alleles are lethal, including null alleles of all subunits we found to change in our proteomics in response to osmotic stress (**Fig. 1D**), but these hypomorphic alleles allow ETC complex function to be partially disrupted without loss of viability (**Fig. 2A**). In addition, MEV-1 is one of the subunits we found to change in abundance in oocytes in response to osmotic stress via proteomics (**Fig. 1D**).

**Figure 2.**
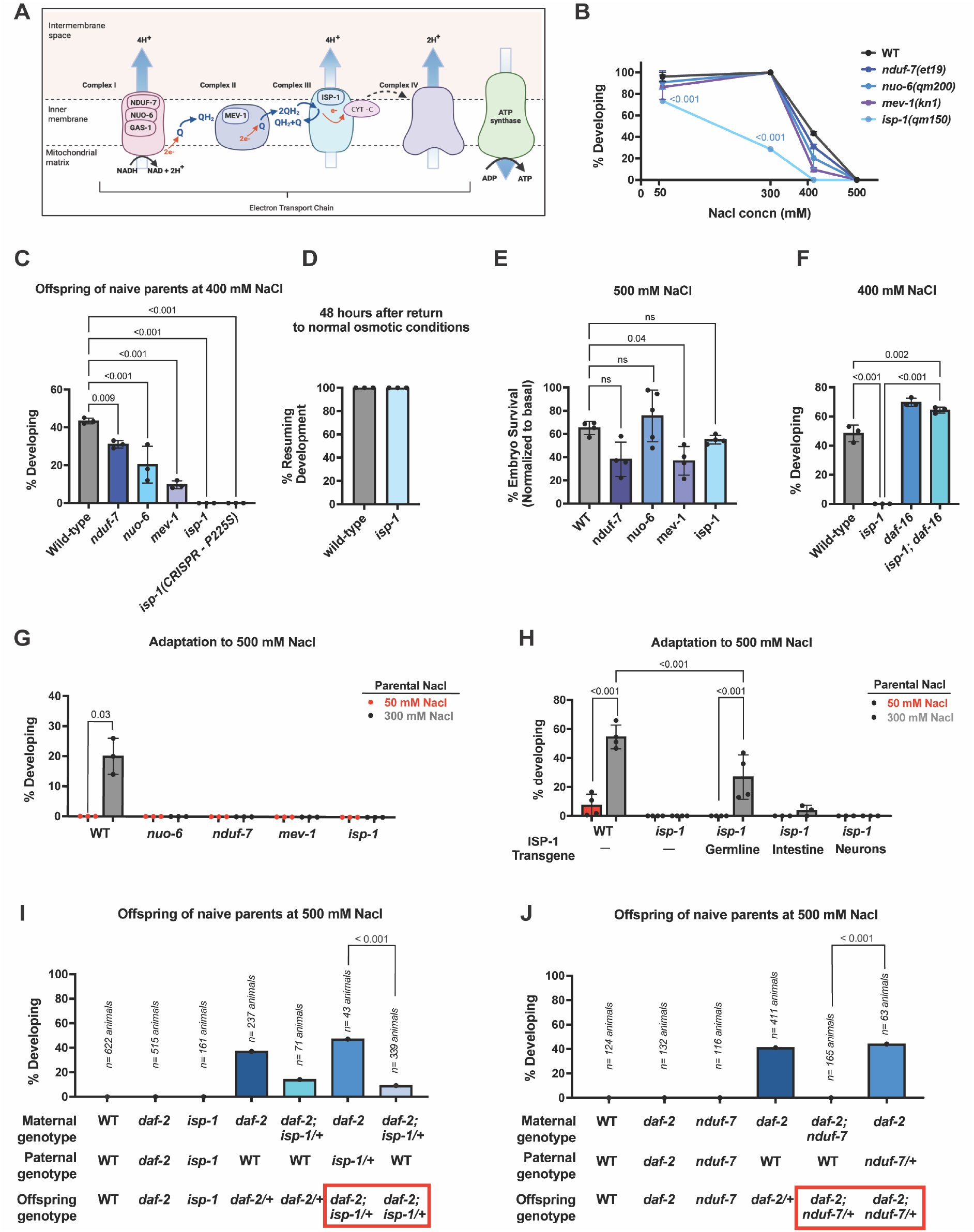
ETC Complex III function regulates animals’ response to osmotic stress. **A.** Schematic showing *C. elegans* Electron transport chain (ETC) with existing partial loss-of-function mutations in *C. elegans* are highlighted for each complex in which they exist. **B.** Percent of WT, *nuo-6(qm200), nduf-7(et19), mev-1(kn1),* and *isp-1(qm150)* embryos developing after 48 hours of exposure to 50 mM, 300 mM, 400mM, and 500mM NaCl. n = 3 replicates of > 40 offspring per replicate, Two-way ANOVA with Šídák’s multiple comparisons test, error bars – s.d. **C.** Percent of WT, *nuo-6(qm200), nduf-7(et19), mev-1(kn1), isp-1(qm150),* CRISPR re-created *isp-1(p225S)* embryos developing after 48 hours of exposure to 400 mM NaCl. n = 3 replicates of > 40 offspring per replicate. One-way ANOVA, error bars – s.d. **D.** Percent of WT and *isp-1(qm150)* mutant embryos that arrested at 400 mM NaCl (see Fig. 1C) that were able to resume development upon return to 50 mM NaCl for 48 hours. one-way ANOVA, error bars – s.d. **E.** Percent of WT, *nduf-7, nuo-6, mev-1* and *isp-1* mutant embryos that survived after arresting on 500 mM NaCl for 48 hours and were recovered on 50 mM NaCl for 48 hours. The embryo survival at 500 mM NaCl is normalized to embryo survival at standard 50 mM NaCl conditions for each strain respectively. n = 4 replicates of > 80 embryos one-way ANOVA, error bars – s.d. **F.** Percent of WT, *isp-1(qm150), daf-16(m26)* and *isp-1qm150); daf-16(m26)* mutant embryos that were mobile and developing after 48 hours of exposure to 400 mM NaCl. n = 3 replicates of > 40 offspring per replicate one-way ANOVA, error bars – s.d. **G.** Percent of WT, *nuo-6(qm200), nduf-7(et19), mev-1(kn1),* and *isp-1(qm150)* embryos developing after 48 hours of exposure to 500 mM NaCl after maternal exposure to 50 mM or 300mM NaCl. n = 3 replicates of > 40 offspring per replicate, Two-way ANOVA with Šídák’s multiple comparisons test, error bars – s.d. **H.** Percent of WT, *isp-1(qm150), isp-1(qm150); pie-1p::isp-1*(Germline ISP-1 rescue) *,isp-1(qm150); ges-1p::isp-1* (Intestinal ISP-1 rescue) and *isp-1(qm150); rgef-1p::isp-1* (Neuronal ISP-1 rescue) embryos developing after 48 hours of exposure to 500 mM NaCl after maternal exposure to 50 mM or 300mM NaCl. n = 4 replicates of > 40 offspring per replicate, Two-way ANOVA with Šídák’s multiple comparisons test, error bars – s.d. **I.** Percent of wild-type, *daf-2(e1370),* and *isp-1(qm150)* mutant embryos that develop and are mobile after 48 hours of exposure to 500 mM NaCl. *daf-2(e1370); isp-1(qm150)/+* animals harbor the *nT1[qIs51]* balancer to maintain heterozygous animals. Cross progeny from heterozygous parents were individually genotyped for each animal, and total animals for each cross are listed. Paternal genotype also contained *him-8(e1489); otIs181* to generate male animals, and confirm cross progeny as previously described [**Burton et al**]. The red box highlights genetically identical offspring that have different responses to osmotic stress. Percent of wild-type, *daf-2(e1370),* and *nduf-7(et19)* mutant animals developing and mobile after 48 hours of exposure to 500 mM NaCl. Cross progeny from heterozygous parents were individually genotyped for each animal, and total animals for each cross are listed. Paternal genotype also contained *him-8(e1489); otIs181* to generate male animals and confirm cross progeny as previously described [**Burton et al**]. The red box highlights genetically identical offspring that have different responses to osmotic stress. Data analyzed using logistic regression with Firth’s bias correction. **J.** Percent of wild-type, *daf-2(e1370),* and *nduf-7(et19)* mutant embryos that develop and are mobile after 48 hours of exposure to 500 mM NaCl. Cross progeny from heterozygous parents were individually genotyped for each animal, and total animals for each cross are listed. Paternal genotype also contained *him-8(e1489); otIs181* to generate male animals and confirm cross progeny. The red box highlights genetically identical offspring that have different responses to osmotic stress. Data analyzed using logistic regression with Firth’s bias correction. Source data are provided in Supplemental table 8.

To determine how each of these ETC Complex disruptions affect animals’ response to osmotic stress we first exposed each mutant to a range of osmotic stress conditions (300 mM-500 mM NaCl) to establish baseline response. (Note, historical *gas-1(fc21)*^30^, Complex I mutants were not used in this assay due to a unique sensitivity of this mutant to egg prep solutions – **Supplementary Fig 2A**). At 300 mM NaCl, we found that 100% of wild-type, *nduf-7*, *mev-1* and *nuo-6* embryos developed normally. Surprisingly, however, the majority of *isp-1* mutant embryos entered the developmental arrest state that occurs specifically in response to osmotic stress (**Fig. 2B**). (We note here that *C. elegans* arrest in response to osmotic stress is a unique regulated arrest response that is distinct from L1 arrest in response to starvation, and is phenotypically similar to the suspended animation states many invertebrates enter in response to osmotic stress – See Burton et al. 2017 for further details^3^). At 400 mM NaCl, which causes a substantial portion of wild-type animals to arrest their development, 100% of *isp-1* mutants were developmentally arrested (**Fig. 2B, C**). By contrast, *nduf-7, mev-1*, and *nuo-6* mutants exhibited only a small, but statistically significant, increase in the percentage of animals entering the arrest state (**Fig. 2B, C**). Lastly, we confirmed our findings for *isp-1* mutants were due to the *isp-1* P225S mutation by recreating it with CRISPR/Cas9 (**Fig. 2C**). This arrest was reversible, as animals returned to normal osmotic conditions resumed development (**Fig. 2D**), confirming that *isp-1* mutants were entering a developmental arrest state rather than simply dying due to overwhelming stress. This suggests that while all the tested ETC subunits seem important for resilience against osmotic stress, only the Complex III subunit (*isp-1*) conferred a unique preference to arrest development in response to mild osmotic stress.

The increased sensitivity of *isp-1* mutants to osmotic stress could reflect a general frailty caused by ETC dysfunction or an active regulatory role for Complex III in the decision to enter a stress resistant state of developmental arrest under stress. We distinguished between these possibilities in two ways. First, we hypothesized that a general frailty should also lead to increased lethality in response to severe osmotic stress. However, we found that *nduf-7*, *nuo-6*, and *isp-1* mutant embryos all had similar or better embryo survival at 500mM and 1M NaCl when compared to wild-type animals (**Fig. 2E and Supplementary Fig. 2B**). *mev-1* mutant embryos showed slightly reduced survival at 500mM, but surprisingly, exhibited a similar survival as wild-type animals at 1 M salt conditions (**Fig. 2E**, **Supplementary Fig. 2B**). This finding is consistent with previous studies that also found that *isp-1* mutant adults are more resistant to osmotic stress than wild-type animals^31^. We conclude that these ETC mutants do not exhibit substantially increased lethality in response to osmotic stress.

Second, we hypothesized that an active regulatory role for the ETC (as opposed to frailty) predicts that the loss of a downstream effector of the arrest should restore development even under conditions that cause *isp-1* single mutants to arrest. By contrast, if *isp-1* mutants are generally frail, then the loss of additional proteins that promote animals’ resistance to osmotic stress should further enhance arrest and/or result in lethality. To test this, we crossed *isp-1* mutants to animals with a dysfunctional *daf-16(m26)* allele. *daf-16* encodes the FOXO transcription factor that promotes *C. elegans* stress-induced developmental arrest^3^. We found that *isp-1;daf-16* double mutants developed to adulthood similarly to wild-type animals even at 400 mM NaCl, a condition where all *isp-1* single mutants enter arrest (**Fig 2F**). We conclude that *isp-1* mutants do not simply fail to tolerate osmotic stress, but rather they preferentially and reversibly arrest their development via a DAF-16 dependent mechanism. (*i.e. isp-1;daf-16* double mutants still have a dysfunctional ISP-1, but no longer arrest their development). These findings indicate that Complex III function likely actively regulates this developmental response to osmotic stress via a DAF-16 dependent mechanism, similar to previous findings for other genes that antagonize *C. elegans* arrest in response to osmotic stress (*daf-2, fax-1, unc-42*)^3,32^.

Having established that ETC function shapes an animals’ own sensitivity to osmotic stress, we next asked whether ETC function is also important for the intergenerational adaption to osmotic stress in which maternal exposure to mild stress improves an offspring’s ability to withstand a subsequent, more severe stress. To test this, we exposed wild-type and ETC mutant (*nuo-6, nduf-7, mev-1* and *isp-1)* young adult animals to either 50 mM (control) or 300 mM NaCl and then assayed the development of their offspring when exposed to 500 mM NaCl. As previously reported, all wild-type offspring of 50 mM-exposed parents arrested their development at 500 mM NaCl, while >20% of offspring from 300 mM-exposed parents developed successfully at 500 mM NaCl (**Fig. 2G**). By contrast, we found that 100% of the ETC mutant offspring arrested at 500 mM NaCl regardless of parental exposure (**Fig. 2G**), demonstrating that ETC function is required for intergenerational adaptation. To test if these partial loss-of-function mutants retained some ability to adapt to mild osmotic stress (consistent with their retention of partial protein function) we also assayed the response of ETC mutants to the mild stress of 400 mM NaCl with and without parental exposure to mild stress (300 mM NaCl). We found that *nduf-7, nuo-6* and *mev-1* mutants retained partial adaptive capacity to 400 mM NaCl, while *isp-1* mutants remained 100% arrested (**Supplemental Fig. 2C**), consistent with its increased preference to arrest embryo development in response to mild hyperosmotic stress in naïve animals (**Fig. 2B**). These findings indicate that while ETC mutants exhibit only a small defect in the response of naïve animals to osmotic stress (**Figs. 2B-C**) these same mutants are completely defective in adapting to 500 mM NaCl (**Fig. 2G**).

*C. elegans* heritable adaptation to osmotic stress involves maternal somatic tissue, maternal oocytes, and offspring somatic tissue^3^. ETC function in any of these tissues or generations could impact this adaptation. Based on the oocyte ETC proteome remodeling we observed in response to osmotic stress (**Fig 1D**), we hypothesized that ETC function specifically in oocytes promotes this heritable adaptation. To test this, we generated transgenic lines expressing a rescue copy of *isp-1(+)* specifically in germ cells (*pie-1* promoter), neuronal cells (*rgef-1* promoter) or intestinal cells (*ges-1* promoter). These tissues were chosen because they have all been previously linked to *C. elegans* heritable adaptation to osmotic stress^3^, and we validated robust expression of the ISP-1 transgene across lines (**Supplemental Fig. 2D)**. Using these lines we found that germline, but not neuronal or intestinal expression of *isp-1(+)* partially rescued both *isp-1(qm150)* mutants’ sensitivity to 400 mM NaCl (**Supplemental Fig. 2E**) and their ability to heritably adapt to 500 mM NaCl (**Fig. 2H)**. A weaker, partial rescue from intestinal *isp-1*(+) expression was observed only at a milder stress level (450 mM NaCl) (**Supplemental Fig 2F)**. Because germline-rescued animals still lack functional *isp-1* in somatic cells, this germline expression rescue further argues against a model in which somatic ETC dysfunction alone drives the phenotype (due to frailty). Nonetheless, the mild impact of intestinal rescue suggests that somatic ISP-1 function may make some contribution to the overall adaptation. We conclude that ETC function in germ cells/oocytes is the primary driver of *isp-1* mutants impaired intergenerational adaptation to osmotic stress.

Because transgenic rescue cannot fully exclude the possibility of leaky *isp-1(+)* in non-germ cells we used an independent genetic cross approach to determine if *isp-1* functions maternally. This approach builds on our previous finding that reduced maternal insulin signaling to oocytes is itself sufficient to trigger offspring resistance to future osmotic stress, even in the absence of parental osmotic stress^3^. Briefly, *daf-2* (insulin receptor) mutant mothers, whose reduced insulin signaling to intestinal cells causes them to arrest at even mild osmotic stress, produce offspring that develop normally at 500 mM NaCl when crossed to wild-type males. This initially unexpected effect of reduced maternal insulin signaling becomes observable because the paternal wild-type copy of *daf-2(+)* restores normal insulin signaling to intestinal cells in offspring^3^ (See Burton et al., 2017 for complete details). In other words, this crossing paradigm reveals a maternal effect of reduced insulin signaling to oocytes on offspring stress resistance that is independent of offspring’s own genotype. We reasoned that the same crossing paradigm could test whether *isp-1* is required maternally to promote offspring resistance to osmotic stress. To test this we first attempted to generate *daf-2(e1370);isp-1(qm150)* homozygous double mutant animals, however, we found that these double mutants were 100% sterile. As an alternative approach, we constructed *daf-2(e1370); isp-1(qm150/+)* animals that were heterozygous for the *isp-1(qm150)* mutation and crossed them with wild-type males. We found that offspring from this cross using heterozygous *isp-1(qm150)* mutant mothers mostly arrested their development at 500 mM NaCl regardless of whether they inherited the *isp-1(qm150)* allele or not (**Fig. 2I – columns 5, 7**). By contrast, genetically identical offspring that inherited a copy of the *isp-1(qm150)* mutation from their fathers showed no defect in adapting to 500 mM NaCl (**Fig. 2I – column 6**). We conclude that the function of maternally expressed *isp-1* regulates offspring response to osmotic stress.

To test if disruptions in maternal Complex I function were also sufficient to alter offspring’s response to osmotic stress we performed the same series of crosses as in (**Fig. 2I)** for *daf-2* and *nduf-7* double mutant animals. We found that *daf-2;nduf-7* double mutant animals were fertile (unlike *daf-2;isp-1* double mutants) and that the loss of maternal, but not paternal, *nduf-7* again altered offspring’s response to 500 mM NaCl independently of offspring genotype (**Fig. 2J**). Therefore, we conclude that both maternal Complex I and Complex III function are required for reduced maternal insulin signaling to promote offspring resistance to future osmotic stress, consistent with our previous tissue-specific rescue results (**Fig. 2H**).

### Oocyte glycogen remodeling in response to osmotic stress promotes intergenerational stress adaptation

Studies in fruit fly and frog oocytes have reported that oocyte ETC function and proteome remodeling are fundamentally linked to glycogen metabolism^17^. Furthermore, original studies of *C. elegans* intergenerational adaptation to osmotic stress noted that this adaptation correlated with reduced glycogen abundance embryos, although the functional significance of these glycogen changes and if they might impact the observed adaptation was unknown^33^. Based on these findings, we hypothesized that oocyte ETC function might regulate an offspring’s response to future osmotic stress by altering oocyte glycogen abundance and/or its metabolism. To test this, we first asked whether maternal exposure to osmotic stress alters glycogen stores in *C. elegans* oocytes. Wild-type animals were exposed to either standard culture conditions (50 mM NaCl) or 300 mM NaCl for 1 hour, 6 hours, or 24 hours and glycogen abundance was quantified by Lugol’s staining (**Fig 3A-C**). We observed high amounts of glycogen (in pre-fertilized -1, -2, and -3) oocytes of wild-type animals grown under normal osmotic conditions (**Fig. 3A-C and Supplemental Fig. 3A)**. This is consistent with previous work showing high levels of glycogen within oocytes right before fertilization in other species^17^. By contrast, exposure of animals to mild salt stress (300 mM NaCl) led to the depletion of these glycogen reserves in as little as 1 hour and this depletion persisted following 24 hours of salt exposure (**Fig. 3A-C and Supplemental Fig. 3A)**. Thus, maternal exposure to osmotic stress leads to significantly reduced glycogen stores in oocytes.

**Figure 3:**
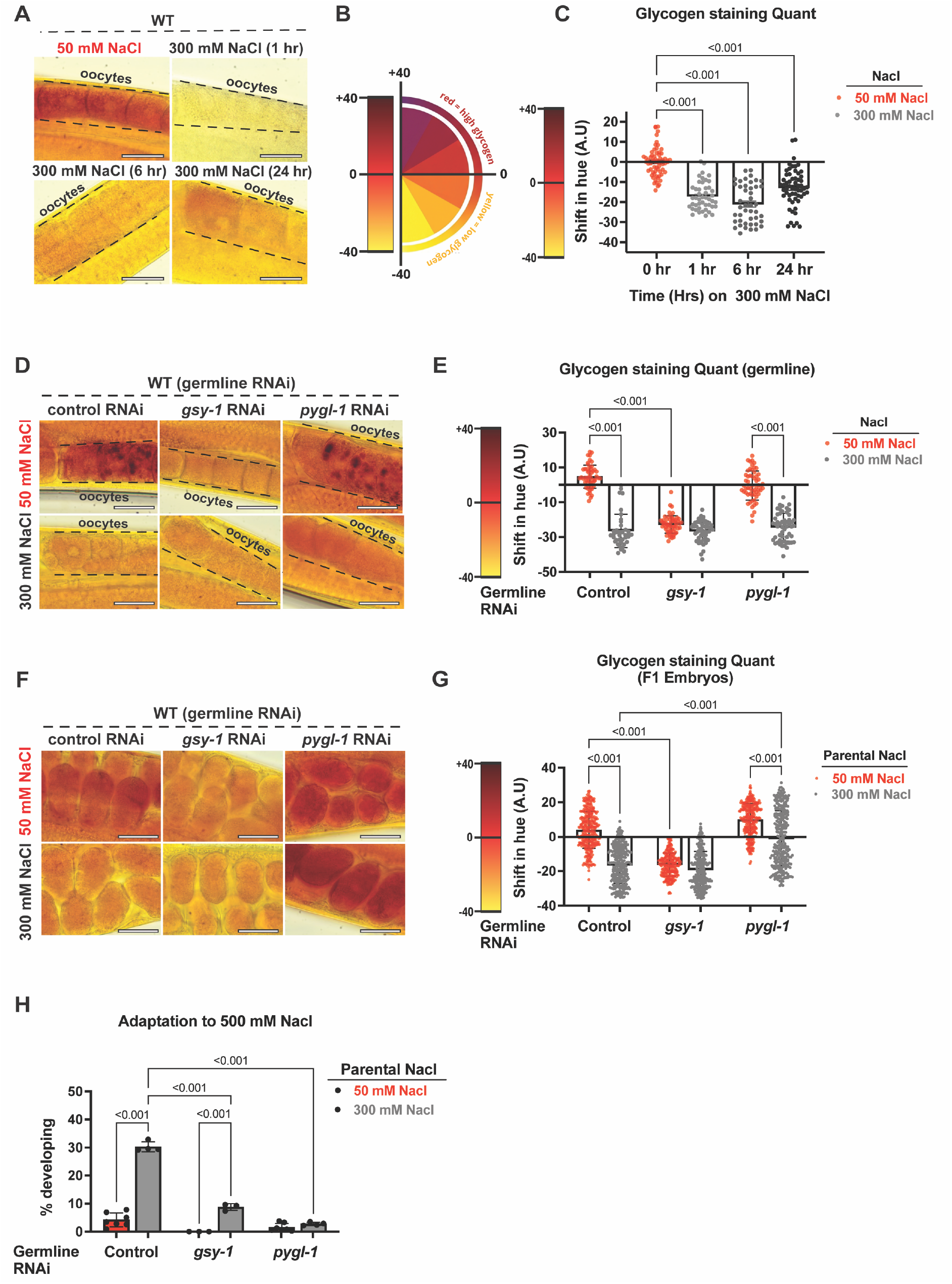
Maternal oocyte glycogen metabolism regulates offspring’s response to future osmotic stress. **A.** Representative brightfield images of lugol’s iodine-stained (reveals glycogen levels) oocytes of day-1 adult WT worms at standard conditions (50 mM NaCl) and after different time points (1, 6 and 24 hours) following mild salt stress (300 mM Nacl). **B.** Representative colour wheel showing different hues of oocyte glycogen observed after lugol’s iodine staining. **C.** Quantification of Fig. 3A. Shift in hue is calculated for each condition with respect to standard condition (50 mM NaCl). n=3 replicates, 20 animals imaged per replicate, one-way ANOVA, error bars – s.d. **D.** Representative brightfield images of lugol’s iodine-stained oocytes of *sun-1p::rde-1; rde-1*(*mkc36*) (Germline specific-RNAi) mothers. Worms were grown on control, *gsy-1* or *pygl-1* RNAi at standard conditions (50 mM NaCl) and at early day-1 of adulthood either exposed to 50 mM or mild salt stress (300 mM Nacl) for 24 hours followed by lugol’s staining. **E.** Quantification of Fig. 3D. Shift in hue is calculated with respect to worms grown on control RNAi at 50 mM NaCl. n=3 replicates, 20 animals imaged per replicate, one-way ANOVA, error bars – s.d. **F.** Representative brightfield images of lugol’s iodine-stained embryos from *sun-1p::rde-1;rde-1*(*mkc36*) (Germline specific-RNAi) worms. Worms were grown on control, *gsy-1* or *pygl-1* RNAi at standard conditions (50 mM NaCl) and either exposed to 50 mM or mild salt stress (300 mM Nacl) for 24 hours followed by lugol’s staining of respective F1 embryos in uterus of these mothers. **G.** Quantification of Fig. 3F. Shift in hue is calculated with respect to worms grown on control RNAi at 50 mM NaCl. n = 4 replicates, 20 animals imaged per replicate, one-way ANOVA, error bars – s.d. **H.** Percent of embryos developing at 500 mM NaCl. *sun-1p::rde-1;rde-1*(*mkc36*) (Germline specific-RNAi) worms were grown on control, *gsy-1* or *pygl-1* RNAi at standard conditions (50 mM NaCl) and at early day-1 of adulthood were either exposed to 50 mM or mild salt stress (300 mM Nacl) for 24 hours. The embryos from these mothers were exposed to 500 mM NaCl for 48 hours, following which developing and mobile larva were scored. n = 4 replicates of > 80 offspring per replicate, Two-way ANOVA with Tukey’s multiple comparison test. error bars -s.d. Scale bars: 30 μM. Source data are provided in Supplemental table 8.

We next asked whether glycogen metabolism in oocytes was functionally required for intergenerational adaptation to osmotic stress. To test this, we performed RNAi silencing of *gsy-1* (glycogen synthase)^33,34^ and the glycogen breakdown gene *pygl-1* (glycogen phosphorylase)^35,36^ specifically in germ cells using a *C. elegans* line that is only capable of RNAi silencing in germ cells^37^. As expected, germline RNAi of *gsy-1* led to a marked loss of glycogen stores in both maternal oocytes and resulting embryos (**Fig 3D-G and Supplemental Fig. 3B, C)**, consistent with a defect in glycogen synthesis. By contrast, *pygl-1* RNAi in germ cells led to glycogen accumulation in embryos but not in maternal oocytes **(Fig 3D-F and Supplemental Fig. 3B, C)**. This is consistent with PYGL-1’s known function in glycogen breakdown and suggests that oocyte derived PYGL-1 protein (or mRNA) is required for glycogen breakdown in embryos. We confirmed that *pygl-1* RNAi results in an ∼95% reduction in *pygl-1* mRNA in dissected oocytes without affecting *pygl-1* mRNA abundance in embryos and that the effects on embryo glycogen abundance persisted even in a *wago-3* mutant background^38^, indicating that the phenotype was not due to inherited RNAi silencing (**Supplemental Fig. 3D-F**).

We next tested whether these perturbations of glycogen metabolism affected *C. elegans* intergenerational adaptation to osmotic stress. Neither RNAi of *gsy-1* nor *pygl-1* in germ cells significantly altered embryonic viability under standard growth conditions (**Supplemental Fig. 3G**). However, when parents were exposed to mild osmotic stress (300 mM NaCl) RNAi silencing of *gsy-1* or *pygl-1* within the germline blocked offspring’s ability to adapt to future osmotic stress (**Fig 3H and Supplemental Fig. 3H).** We conclude that both glycogen synthesis and breakdown in maternal germ cells/early F1 embryos are required for animals’ ability to heritably adapt to osmotic stress.

Animals that fail to adapt to osmotic stress could fail to adapt for several reasons. First, animals could fail to adapt because they either preferentially arrest their development in response to osmotic stress or because they cannot exit developmental arrest. In this case, all animals remain alive and often exhibit enhanced survival in response to severe osmotic stress as we observed here for *isp-1* mutants (**Fig 2** and **Supplemental Fig 2B**) and previously for *daf-2* mutants^3^. By contrast, animals that are unable to respond properly to osmotic stress could fail to adapt because osmotic stress becomes lethal. This scenario would be similar to our previous findings for *gpdh-2* mutants which cannot synthesize glycerol (an osmolyte) as part of the response to osmotic stress. To differentiate between these possibilities, we performed RNAi silencing of *gsy-1* or *pygl-1* in germ cells of parents exposed to either 50 mM or 300 mM NaCl and assayed the survival of embryos at 500 mM NaCl. We found that while RNAi of *gsy-1* or *pygl-1* in germ cells did not affect survival at normal osmotic conditions (**Supplemental Fig 3G**), it led to increased lethality when embryos were exposed to 500 mM NaCl (**Supplemental Fig 3I**). These findings are consistent with a model in which the metabolism of oocyte derived glycogen is required to properly respond to osmotic stress, potentially by fueling the production of osmolytes similarly to the function of GPDH-2 in the osmotic stress response.

### Maternal insulin signaling and oocyte ETC function both regulate oocyte glycogen abundance

We next asked whether two maternal pathways previously linked to this intergenerational adaptation -insulin signaling and oocyte ETC function (**Fig 2**) -might both regulate oocyte glycogen metabolism. Previous studies of fly oocytes found that reduced insulin signaling led to increased glycogen abundance^17^. Consistent with these findings, we found that *daf-2* mutant *C. elegans* also accumulated substantially increased glycogen abundance in oocytes when compared to wild-type animals, and this accumulation required glycogen synthase (*gsy-1*) in oocytes (**Fig. 4 A,B, Supplemental Fig. 4A**). Thus, maternal insulin signaling antagonizes glycogen synthesis in oocytes. Because loss of maternal insulin signaling to oocytes is sufficient to drive offspring resistance to future osmotic stress even the absence of maternal osmotic stress^3^, we hypothesized that increased glycogen accumulation in *daf-2(e1370)* mutant oocytes might contribute to this phenotype. As above, germline RNAi silencing of *gsy-1* prevented glycogen accumulation in *daf-2* mutant animals (**Fig 4 C, D Supplemental Fig. 4B**). Thus, to test our hypothesis, we crossed *daf-2(e1370)* mutant mothers +/-germline specific RNAi of *gsy-1* and *pygl-1* with wild-type males and assayed their offspring’s response to strong osmotic stress (500 mM NaCl). Although loss of maternal insulin signaling (followed by inheritance of a wild-type copy of *daf-2* from their fathers) normally protected offspring against developmental arrest, RNAi of either *gsy-1* or *pygl-1* in germ cells significantly reduced this protective effect (**Fig 4 E, F**). These findings suggest that reduced maternal insulin signaling protects offspring against future osmotic stress via a mechanism that, at least in part, requires increased glycogen accumulation in oocytes and breakdown of oocyte-derived glycogen via PYGL-1 in offspring.

**Figure 4:**
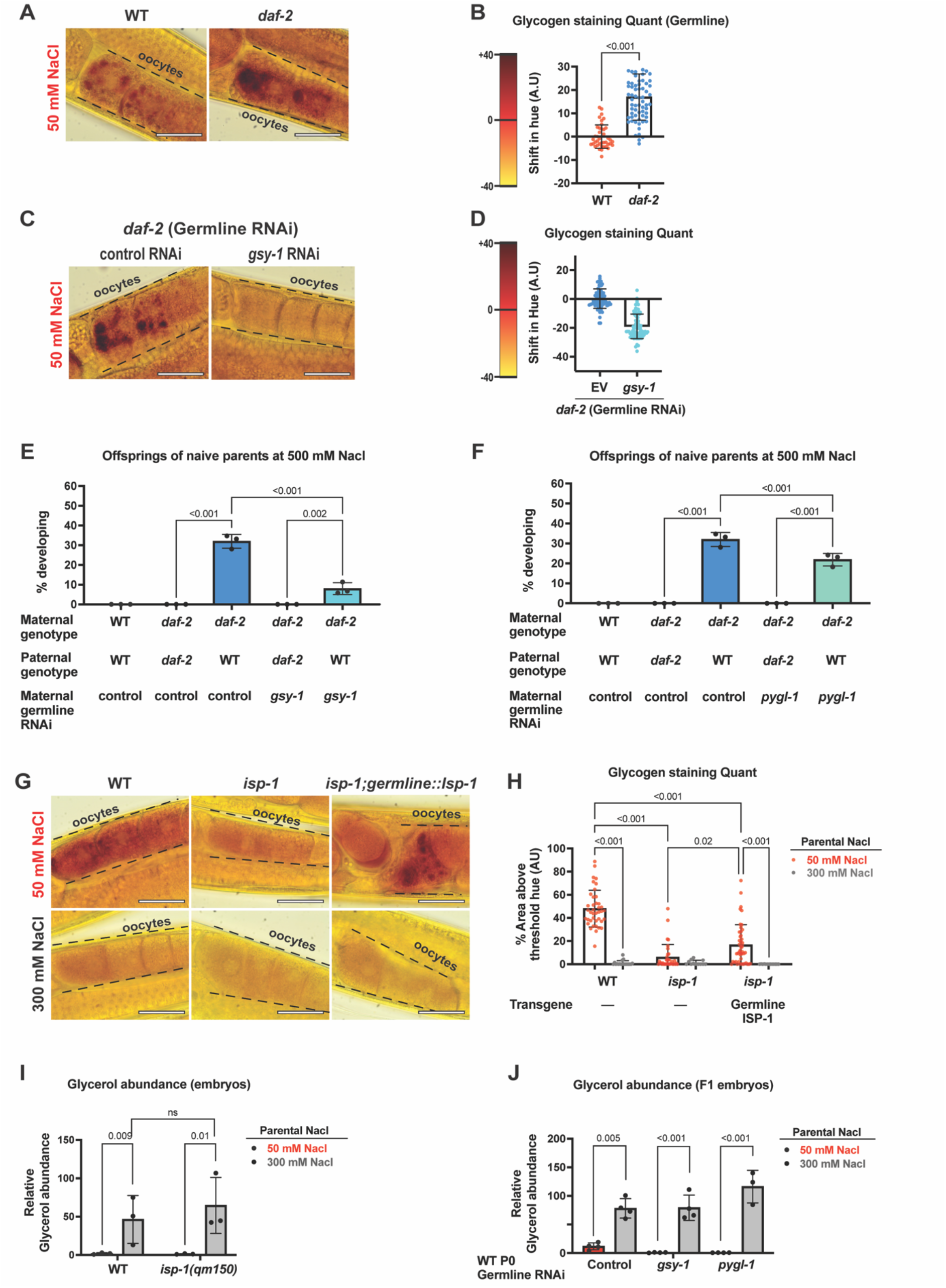
Insulin signalling and mitochondrial ETC antagonistically influence maternal oocyte glycogen levels. **A.** Representative brightfield images of lugol’s iodine-stained oocytes of *daf-2* and WT day-1 adults at standard conditions (50 mM NaCl). **B.** Quantification of Fig. 4A. Shift in hue is calculated with respect to WT. n=3 replicates, 20 animals imaged per replicate, Unpaired students’ *t*-test with Welch correction, error bars – s.d. **C.** Representative brightfield images of lugol’s iodine-stained oocytes of day-1 adult *daf-2(e1370); sun-1p::rde-1 ;rde-1(mkc36)* (Germline specific-RNAi) worms grown on control or *gsy-1* RNAi at standard conditions (50 mM NaCl). **D.** Quantification of Fig. 4C. Shift in hue is calculated with respect to *daf-2(e1370)* worms grown on control RNAi. n=3 replicates, 20 animals imaged per replicate, Unpaired students’ *t*-test with Welch correction, error bars – s.d. **E.** Percent of embryos developing at 500 mM NaCl. *daf-2(e1370); sun-1p::rde-1; rde-1(mkc36)* worms were grown on control or *gsy-1* RNAi at standard conditions (50 mM NaCl) and crossed to WT or *daf-2(e1370)* males grown on control RNAi. Simultaneously, WT and *daf-2(e1370); sun-1p::rde-1; rde-1(mkc36)* worms grown on control RNAi were self-mated. The F1 embryos from self-mating and cross mating were exposed to 500 mM NaCl for 48 hours, after which developing and mobile larva were transferred to OP50 seeded plates and scored 48 hours later to ascertain cross-progeny that developed. n= 3 replicates, 40 animals per replicate, one-way ANOVA, error bars – s.d. **F.** Percent of embryos developing at 500 mM NaCl. *daf-2(e1370) ;sun-1p::rde-1; rde-1(mkc36)* worms were grown on control or *pygl-1* RNAi at standard conditions (50 mM NaCl) and crossed to WT or *daf-2(e1370)* males grown on control RNAi. Simultaneously, WT and *daf-2(e1370); sun-1p::rde-1; rde-1(mkc36)* worms grown on control RNAi were self-mated. The F1 embryos from self-mating and cross mating were exposed to 500 mM NaCl for 48 hours, after which developing and mobile larva were transferred to OP50 seeded plates and scored 48 hours later to ascertain cross-progeny that developed. n= 3 replicates, 40 animals per replicate, one-way ANOVA, error bars – s.d. **G.** Representative brightfield images of lugol’s iodine-stained oocytes of WT, *isp-(qm150)* and *isp-(qm150); pie-1p::isp-1*(Germline *isp-1* rescue) adult worms. Worms were grown at standard conditions (50 mM NaCl) and at early day-1 of adulthood either exposed to 50 mM or mild salt stress (300 mM Nacl) for 24 hours followed by lugol’s staining. **H.** Quantification of Fig. 4G. Shift in hue is calculated for each condition with respect to WT on standard condition (50 mM NaCl). n=3 replicates, >12 animals imaged per replicate, one-way ANOVA, error bars – s.d. **I.** Relative glycerol abundance in embryos. WT and *isp-(qm150)* worms were grown at standard conditions (50 mM NaCl) and at early day-1 of adulthood either exposed to 50 mM or mild salt stress (300 mM Nacl) for 24 hours. Following this, embryos were isolated from each condition and processed utilizing gas chromatography mass spectroscopy (GC-MS) to assess relative glycerol abundance. n= 3 replicates, Two-way ANOVA with Šídák’s multiple comparisons test, error bars – s.d. **J.** Relative glycerol abundance in embryos. *sun-1p::rde-1; rde-1*(*mkc36*) (Germline specific-RNAi) worms were grown on control, *gsy-1* or *pygl-1* RNAi at standard conditions (50 mM NaCl) and at early day-1 of adulthood were either exposed to 50 mM or mild salt stress (300 mM Nacl) for 24 hours. Following this, embryos were isolated from each condition and processed utilizing gas chromatography mass spectroscopy (GC-MS) to assess relative glycerol abundance. n= 3 or 4 replicates, Two-way ANOVA with Šídák’s multiple comparisons test, error bars – s.d. Scale bars: 30 μM. Source data are provided in Supplemental Table 8.

We next asked whether the oocyte ETC function, identified as involved in the adaptation above (**Fig 2**), also impacted oocyte glycogen abundance. Using our *isp-1* mutant model, we found that the loss of ISP-1 function led to decreased glycogen accumulation in oocytes (**Fig 4 G, H**). Furthermore, we found that germline specific expression of a wild-type copy of *isp-1(+)* was sufficient to increase glycogen abundance in a fraction of our otherwise *isp-1* mutant animals (**Fig 4 G, H**). A similar reduction in oocyte glycogen was also observed for *nduf-7* (Complex I mutant) animals (**Supplemental Fig. 4 C-E**). These findings indicate that in *C. elegans,* ETC function in oocytes promotes glycogen accumulation in oocytes. Notably, previous studies of fly oocytes proposed that ETC remodeling led to reduced respiratory activity, which in turn was hypothesized to promote increased oocyte glycogen accumulation (*i.e.* that oocyte ETC function and oocyte glycogen stores were inversely related)^17^. However, this model had not been directly tested by perturbing ETC function in oocytes. Our findings suggest that oocyte ETC function, at least in *C. elegans*, promotes increased oocyte glycogen abundance.

Taken together, our findings indicate that maternal insulin signaling and oocyte ETC function both regulate oocyte glycogen abundance, although they do so in different directions. Reduced insulin signaling increases oocyte glycogen, whereas disruption of ETC function decreases it. In addition, our findings indicate that genetic methods that drive increased glycogen accumulation in oocytes (by reducing maternal insulin signaling) are sufficient to allow offspring to bypass developmental arrest in response to strong osmotic stress (500 mM NaCl) even if the parents themselves never experienced any osmotic stress. By contrast, all of our genetic methods that perturb glycogen accumulation in oocytes or its metabolism (*isp-1* mutants and RNAi of *gsy-1* and *pygl-1* in germ cells) block offspring’s ability to heritably adapt to strong osmotic stress (500 mM NaCl). Collectively, these findings suggest that changes in oocyte glycogen metabolism are both necessary and sufficient to promote offspring’s future resistance to osmotic stress, at least in these contexts.

We have found that maternal exposure to osmotic stress, reduced maternal insulin signaling, and oocyte ETC function all regulate offspring’s future response to osmotic stress (**Fig 2**) and glycogen metabolism in oocytes. (**Figs 3-4**). These findings could be because all three of osmotic stress exposure, maternal insulin signaling, and oocyte ETC activity function as part of a single linear pathway or because all three mechanisms converge on the regulation of oocyte glycogen stores in parallel. To differentiate between these two models, we performed proteomics profiling of wild-type and *daf-2* mutant oocytes under standard osmotic conditions to test if the loss of insulin signaling alone was sufficient to trigger the same changes in ETC protein abundance in oocytes that we observed in response to osmotic stress (**Fig 1D**). This experiment revealed no significant overlap in proteins altered by loss of maternal insulin signaling and those altered by maternal osmotic stress (**Supplemental Fig 4F, Supplemental Table 5**). Thus, the loss of maternal insulin signaling alone was not sufficient to alter ETC protein abundance in *C. elegans* oocytes. In addition to these findings, we note that the loss of maternal insulin signaling alone was sufficient to drive increased glycogen accumulation in oocytes (**Fig 4 A-B**) while maternal exposure to osmotic stress resulted in the metabolism of oocyte glycogen (**Fig 3A-C**). These collective findings support a model in which maternal exposure to osmotic stress and the loss of maternal insulin signaling to oocytes promote offspring’s resistance to osmotic stress via parallel effects on oocyte glycogen. Our data further support a model in which the metabolism of oocyte-derived glycogen (via PYGL-1) drives offspring’s resistance to future stress and this metabolism can occur either in embryos once they are exposed to osmotic stress (which inherit more oocyte derived glycogen from *daf-2* mutant mothers) or it can begin already in oocytes in mothers who are actively experiencing osmotic stress and then continue in embryos. However, we also note that we previously demonstrated that maternal exposure to osmotic stress itself causes reduced insulin signaling too oocytes^3^. Thus, it is likely that maternal osmotic stress exposure and maternal insulin signaling exhibit some level of intersection and crosstalk to regulate offspring’s future response to osmotic stress.

### Maternal ETC function and oocyte glycogen metabolism regulates *C. elegans* response to osmotic stress via an AMPK-kinase dependent mechanism in offspring

The metabolism of oocyte derived glycogen could impact offspring’s future response to osmotic stress by simply serving as a fuel for glycerol production. Alternatively, the metabolism of oocyte-derived glycogen could regulate the activity of downstream embryo signaling pathways that promote future stress resistance. Specifically, we previously found that glycerol production is substantially increased in offspring from parents exposed to osmotic stress and that increased glycerol production is required for *C. elegans* heritable adaptation to osmotic stress. Because glycogen can be broken down to glucose, and in principle this glucose could be utilized to make glycerol, we hypothesized that oocyte glycogen might provide the fuel for increased glycerol accumulation in offspring of stressed parents. Unexpectedly, embryo glycerol levels were not reduced in the offspring of *isp-1* mutant parents exposed to 300 mM NaCl (**Fig 4I**). Similarly, germline RNAi of *gsy-1* or *pygl-1* did not affect glycerol accumulation in offspring when parents were exposed to 300 mM NaCl (**Fig. 4J**). These findings suggest that while defects in ETC function and glycogen synthesis/breakdown block *C. elegans* intergenerational adaptation to osmotic stress, they do so via a mechanism that is independent from glycerol synthesis and accumulation in embryos in response to osmotic stress.

To identify the glycerol-independent downstream mechanisms by which oocyte glycogen metabolism might regulate offspring’s future response to osmotic stress, we conducted two independent mutagenesis screens. One screen was performed on wild-type animals to identify mutants that could bypass developmental arrest when exposed to 500 mM NaCl (this is the same screen that previously identified *ssu-1* and *nhr-1* as genes that regulate animals response to osmotic stress)^39^. The second screen was performed on *isp-1* animals to identify suppressor mutants that could restore development to these ETC mutants under mild and strong osmotic stress conditions (>300 mM NaCl). Interestingly, both screens identified alleles of the AMP kinase (AMPK) catalytic subunit *aak-2* (**Fig. 5A-C, Supplemental Fig. 5A**). An independent *aak-2(ok524)* deletion allele reproduced the phenotypes identified in the screens, and loss of *aak-2* restored the ability of *isp-1* mutants to develop at 350 mM NaCl (**Fig. 5B-C, Supplemental Fig. 5A)**. Thus, loss of AAK-2 suppresses the stress-induced developmental arrest caused by loss of ISP-1, consistent with a model in which ETC function regulates the decision to arrest development through a mechanism that requires AMP-kinase activity. These results suggest that the metabolism of oocyte-derived glycogen could impact offspring resistance to future osmotic stress via altering embryo AMP-kinase signaling.

**Figure 5:**
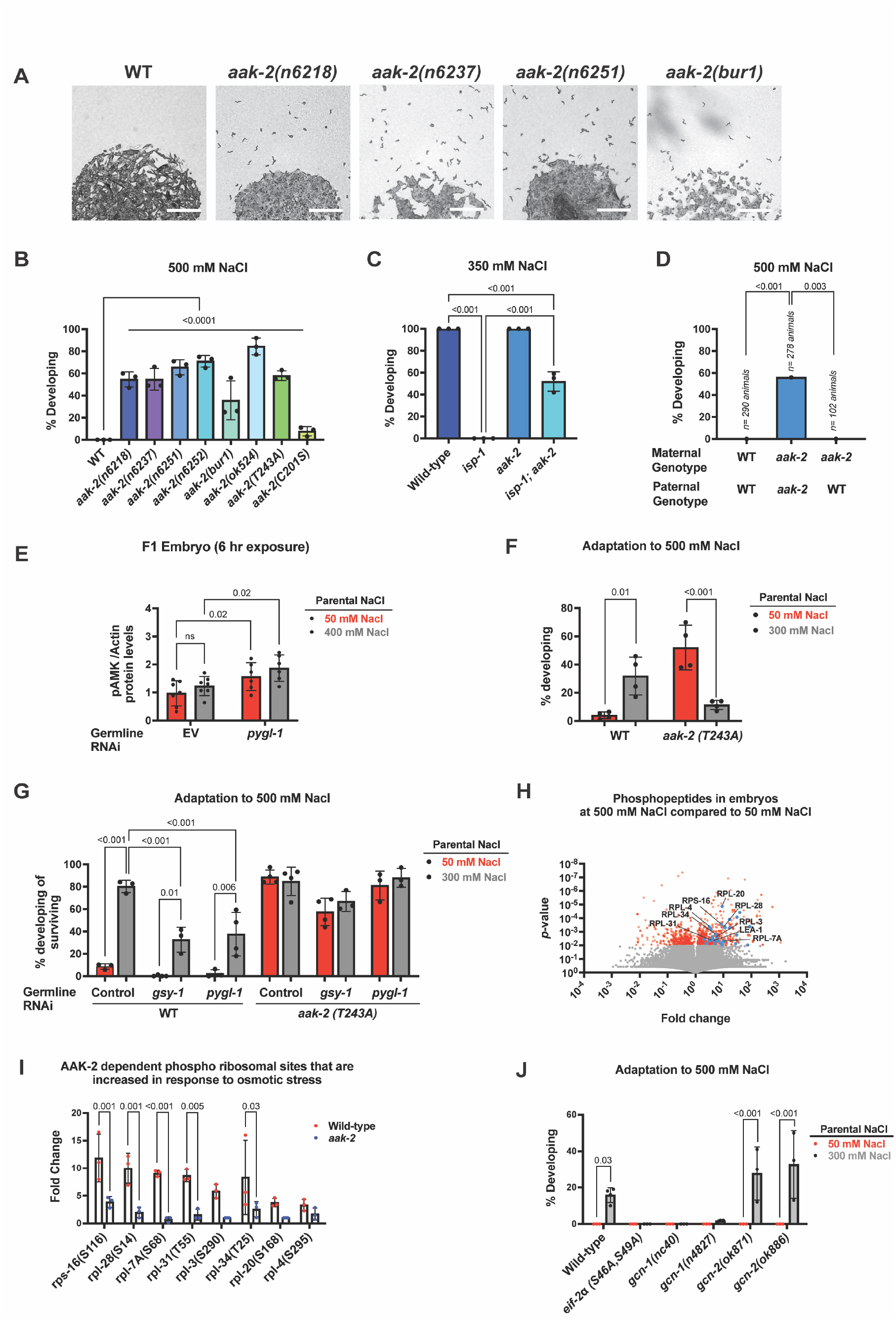
ETC Complex III dysfunction regulates *C. elegans* response to osmotic stress via and AMP-kinase/*aak-2* dependent mechanism. **A.** Representative images of >500 wild-type and *aak-2* mutant *C. elegans* embryos placed on NGM agar plates containing 500 mM NaCl for 48 hours. 100% of wild-type animals enter developmental arrest at hatching, while a fraction of *aak-2* mutants bypass developmental arrest and continue developing. For imaging purposes, plates contained no food and mobile animals remain in the L1 larval stage. Scale bars: 1 mm. **B.** Percent of WT and *aak-2* mutant animals developing and mobile after 48 hours of exposure to 500 mM NaCl. n = 3 replicates of >100 animals per replicate. one-way ANOVA, error bars – s.d. **C.** Percent of WT, *aak-2(n6251)*, *isp-1(qm150)*, and *isp-1*(qm150); *aak-2(n6251)* double mutant embryos that develop and become mobile after 24 hours of exposure to 350 mM NaCl. n = 3 replicates of >100 animals per replicate. one-way ANOVA, error bars – s.d. **D.** Quantification of viable WT and *aak-2(n6251)* mutant cross progeny developing and mobile after 48 hours exposure to 500 mM NaCl cultures. Paternal genotypes also contained *him-8(e1489); otIs181* which generate large numbers of males and also to confirm cross progeny, as previously described in [Burton, N. O. et al. Insulin-like signaling to the maternal germline controls progeny response to osmotic stress. Nat. Cell Biol. 19, 252–257 (2017).] Data analyzed using logistic regression with Firth’s bias correction. **E.** Quantification of western blot as shown in Supplementary Figure 5B. Relative pAMPK levels with respect to actin are plotted. Briefly, *sun-1p::rde-1; rde-1(mkc36)* (Germline specific-RNAi) animals were grown on control or *pygl-1* RNAi till day-1 of adulthood. The embryos were isolated and exposed to 50 mM or 400 mM NaCl for 6 hours. Animals were then collected and embryos extracted for western blot analysis. Actin was probed as a loading control. Two-way ANOVA with Šídák’s multiple comparison test. n = 6, error bars – s.d. **F.** Percent of embryos developing at 500 mM NaCl. *sun-1p::rde-1; rde-1*(*mkc36*) (Germline specific-RNAi) and *aak-2(T243A); sun-1p::rde-1; rde-1*(*mkc36*) worms were grown on control RNAi at standard conditions (50 mM NaCl) and at early day-1 of adulthood were either exposed to 50 mM or mild salt stress (300 mM Nacl) for 24 hours. The embryos from these mothers were exposed to 500 mM NaCl for 48 hours, following which developing and mobile larva was scored and % developing worms were plotted with respect to total number of eggs. n = 4 replicates of > 80 offspring per replicate, Two-way ANOVA with Tukey’s multiple comparison test. error bars -s.d. **G.** Percent of embryos developing at 500 mM NaCl. *sun-1p::rde-1; rde-1*(*mkc36*) (Germline specific-RNAi) and *aak-2(T243A); sun-1p::rde-1; rde-1*(*mkc36*) worms were grown on control, *gsy-1* or *pygl-1* RNAi at standard conditions (50 mM NaCl) and at early day-1 of adulthood were either exposed to 50 mM or mild salt stress (300 mM Nacl) for 24 hours. The embryos from these mothers were exposed to 500 mM NaCl for 48 hours, following which developing and mobile larva was scored and arrested embryos were rescued and scored. % developing worms were plotted with respect to total number of survivors, n = 3 or 4 replicates of > 80 offspring per replicate, Two-way ANOVA with Tukey’s multiple comparison test. error bars -s.d. **H.** Volcano plot of 112,498 detected phospho-peptides that passed all filters in 3 biological replicates of wild-type (WT) embryos collected from day-1 mothers. Embryos exposed to either standard culture conditions (50 mM NaCl) or osmotic stress (500 mM NaCl) for 2 hours. Red dots represent phospho-peptides with abundance, *p* < 0.01. Blue dots represent ribosomal phospho-peptides and also include LEA-1. **I.** Fold change in phospho-peptides of ribosomal subunit proteins that change in WT and *aak-2* mutant embryos exposed to osmotic stress (500 mM NaCl) for 2 hours. n = 3 replicates of, Two-way ANOVA with Šídák’s multiple comparison test. error bars -s.d. Percent of embryos developing at 500 mM NaCl. WT, *eif-2α(S46A, S49A)*, *gcn-1(nc40)*, *gcn-1(n4827)*, *gcn-2(ok871)* and *gcn-2(ok886)* worms were grown on control RNAi at standard conditions (50 mM NaCl) and at early day-1 of adulthood were either exposed to 50 mM or mild salt stress (300 mM Nacl) for 24 hours. The embryos from these mothers were exposed to 500 mM NaCl for 48 hours, following which developing and mobile larva was scored. n = 3 or 4 replicates of > 100 offspring per replicate, Two-way ANOVA with Šídák’s multiple comparison test. error bars -s.d. Source data are provided in Supplemental table 8.

AAK-2 is canonically activated *in vivo* by phosphorylating T243 (equivalent to mammalian AMPK T172)^39–42^ and can be noncanonically activated by oxidation of cysteines by ROS^43^. To test if AAK-2 phosphorylation or cysteine oxidation might be required to regulate animals’ response to osmotic stress, we obtained an existing T243A allele of *aak-2* generated by CRISPR/Cas9^44^ and, separately, generated a C201S allele of *aak-2* based on the conservation of C201 (in *C. elegans)* across species. We found that T243A mutants behaved like *aak-2* null mutants, while C201S mutants behaved like weak partial loss-of-function mutants (**Fig. 5B**). We conclude that the canonical phosphorylation site (T243) is required for AAK-2 to promote *C. elegans* entry into a developmental arrest state during periods of osmotic stress.

To test if *aak-2* functions maternally or in offspring, we crossed *aak-2(ok529)* mutant mothers with wild-type males, and quantified offspring response to 500 mM NaCl. Offspring from this cross entered developmental arrest similar to wild-type animals (**Fig. 5D**). We conclude that unlike our findings for ETC subunits (*isp-1, nduf-7*) (**Fig. 2**) and insulin signaling (*daf-2*)^3^, that AAK-2 functions in offspring to regulate animals’ decision to either arrest development or continue developing during periods of osmotic stress.

Regulation of AMP-kinase activity has already been linked to both ETC function and glycogen abundance in other biological contexts^45,46^. In particular the beta subunit of AMP-kinase contains a glycogen binding domain that can affect both AMP-kinase activation and localization^47^, while glycogen metabolism can influence the ATP/AMP ratio, another well-established regulator of AMP-kinase activation^35,48,49^, by fueling ETC function. These collective observations led us to hypothesize that oocyte glycogen abundance and its subsequent metabolism might directly regulate AAK-2 activity in embryos. If offspring AAK-2 functions in a signaling pathway that is regulated by oocyte glycogen metabolism, then we hypothesized that an inability to metabolize oocyte glycogen would lead to increased phosphorylation of AAK-2 at Thr243 which in turn would increase offspring AAK-2 activity and promote animals’ entry into developmental arrest at 500 mM NaCl. To test this model, we performed RNAi silencing of *pygl-1,* the glycogen metabolism gene that had the strongest effect on the adaptation phenotype (**Fig 3H**), specifically in germ cells using the same germline specific RNAi strain as above and then assayed the phosphorylation of AAK-2 at Thr243 in offspring. Consistent with our model, we found that RNAi of *pygl-1* in germ cells led to increased phosphorylation of AAK-2 at Thr243 in offspring (**Fig. 5E, Supplemental Fig. 5B**). We conclude that the metabolism of oocyte-derived glycogen via germline expressed PYGL-1 antagonizes the phosphorylation of AAK-2 in embryos. Importantly, these findings suggest that the metabolism of oocyte-derived glycogen functionally connects maternal inputs (environmental stress exposure, insulin signaling, oocyte ETC function) to the activity of AMP-kinase in embryos.

To further understand how AAK-2 activity and glycogen metabolism intersect to regulate animals’ response to osmotic stress we next asked how the simultaneous loss of glycogen metabolism and AAK-2 activity affected animals’ response to osmotic stress. Specifically, we first asked whether RNAi knockdown of *gsy-1* and *pygl-1* in our germline-specific RNAi strain affected the ability of *aak-2* mutants to develop at 500 mM NaCl under baseline conditions. In this case, we found that RNAi knockdown of *gsy-1* and *pygl-1* had no effect on *aak-2*(T243A) mutants’ response to osmotic stress (**Supplemental Fig. 5C**). We next asked whether the loss of *aak-2* could suppress the lethality observed in embryos exposed to 500 mM NaCl when *gsy-1* or *pygl-1* is silenced in germ cells (**Supplemental Fig 3I**). Surprisingly, however, we found that when we exposed *aak-2*(T243A) mutant parents to 300 mM NaCl and assayed their offspring’s response to 500 mM NaCl that >80% of *aak-2*(T243A) mutants no longer survived at 500 mM NaCl (**Fig. 5F**). This finding is consistent with a model in which *aak-2* phosphorylation at T243 is required for animals’ normal response to osmotic stress and that the loss of *aak-2* phosphorylation at T243 becomes lethal under conditions of prolonged (parent and offspring) osmotic stress exposure. In addition, we found that RNAi silencing of *gsy-1* and *pygl-1* in germ cells of *aak-2*(T243A) mutant animals did not further enhance the lethality observed in either condition alone suggesting that they might regulate *C. elegans* survival via a shared mechanism (**Supplemental Fig. 5D**). Despite this significant lethality in the adaptation assay across *aak-2* mutants and RNAi of glycogen metabolism genes in germ cells, we were able to assay the ability of the ∼10% of surviving animals to intergenerationally adapt to 500 mM NaCl. We found that the *aak-2*(T243A) mutation completely suppressed the adaptation defect caused by RNAi silencing of *gsy-1* and *pygl-1* in germ cells in surviving animals (**Fig 5G**). Taken together, these results suggest that the metabolism of oocyte-derived glycogen promotes offspring adaptation to osmotic stress at least in part by antagonizing the phosphorylation of AAK-2 at Thr243.

Our data indicate that AAK-2 functions in embryos to promote animals’ entry into an arrested state in response to osmotic stress. The molecular targets of AAK-2 in animals’ response to osmotic stress, however, are unknown. To identify these targets, we performed global phosphoproteomics profiling of wild-type and *aak-2* mutant embryos exposed to either standard conditions (50 mM NaCl) or osmotic stress (500 mM NaCl) for 2 hours. In doing so, we quantified the relative abundance of >100,000 unmodified and modified peptides (**Fig. 5H, Supplemental table 6 and 7**) and identified 34 phosphopeptides that exhibited a >2-fold increase in phosphorylation in response to osmotic stress that was dependent on *aak-2* (p < 0.01) (**Fig 5I, Supplemental table 6 and 7**). These 34 phosphopeptides included 8 ribosomal subunits, representing a highly significant enrichment (**Fig 5I, Supplemental table 6 and 7**).

Deletion mutants of these core ribosomal subunits do not exist and are presumed to be inviable, preventing us from assaying if these genes are required for *C. elegans* adaptation to osmotic stress. Nonetheless, we noted that recent studies of mouse embryonic fibroblasts (MEFs) found that these mammalian cells enter a “hibernation-like” state in response to strong osmotic stress and that entry into this “hibernation-like” is regulated via control of the ribosome by the Integrated Stress Response (ISR) and the phosphorylation of eIF2alpha^50^. This finding is not only phenotypically similar to our findings for *C. elegans* response to osmotic stress but we note that one of the hallmarks of the ISR, ASNS-2, was the second most significantly increased in abundance protein in embryos from parents exposed to osmotic stress when compared to naïve embryos (**Fig. 1F, Supplemental table 4**)^25^. Based on these collective findings, we hypothesized that AAK-2 and the ISR might converge on regulation of the ribosome to regulate *C. elegans* response to osmotic stress. To test this, we first obtained an established *C. elegans* strain in which S46 and S49 of *eif-2alpha* are mutated to alanine using CRISPR/Cas9 to prevent their phosphorylation^51^. We found that this strain was grossly wild type under normal osmotic conditions, but 0% of animals were able to intergenerationally adapt to 500 mM NaCl (**Fig 5J**). These findings indicate that like in MEFs, *eif-2alpha* phosphorylation is required for *C. elegans* to adapt to osmotic stress.

As a parallel test for the role of the ribosome in *C. elegans* response to osmotic stress we also obtained multiple null alleles of both *gcn-1* (a protein that binds colliding ribosomes and is the central regulator of ribosomal responses to stress)^52–55^ and of one of its downstream effector genes, *gcn-2*. We found that both alleles of *gcn-1* were also unable to intergenerationally adapt to 500 mM NaCl (**Fig 5J**), consistent with the model that ribosomal regulation is part of *C. elegans* intergenerational adaptation to osmotic stress. By contrast, neither allele of *gcn-2* was defecting in adapting to osmotic stress. These results suggest that the ribosome stress sensor GCN-1 regulates animals’ response to osmotic stress and that it does so via a mechanism that is likely independent of GCN-2. Collectively, our findings indicate that the ribosome is phosphorylated in embryos in response to osmotic stress via a mechanism that requires AAK-2 and that disrupting core pathways that regulate the ribosome stress responses (EIF-2alpha, GCN-1) functionally blocks animals’ ability to adapt to osmotic stress.

## Discussion

Collectively, our results establish a novel, intergenerational signaling cascade that links maternal somatic conditions (osmotic stress, insulin signaling) to molecular changes in oocytes (reduced ETC subunit abundance, glycogen metabolism), which in turn regulates the phosphorylation of AMP-kinase in offspring and offspring’s future response to stress (See Model, **Fig 6**). Conceptually, these findings represent several important advances in the field. First, these findings demonstrate that the evolutionarily conserved phenomena of oocyte ETC proteome and glycogen remodeling is sensitive to a mother’s environment, and not simply an intrinsic part of gametogenesis. Second, our findings assign a causal, functional role to this process that links a mother’s environment (and maternal insulin signaling states) to offsprings’ ability to adapt to future stress. Lastly, our findings demonstrate that oocyte ETC function and the metabolism of oocyte-derived glycogen functionally regulate the activity of AMP-kinase in embryos, which likely converges with the ISR to regulate the ribosome as part of animals’ response to osmotic stress. Thus, our findings identify a specific “receiver” of signals from changes in oocyte ETC function and glycogen metabolism that can translate these molecular changes into phenotypic changes in offspring. We believe these findings not only advance our understanding of the molecular mechanisms by which a mother’s environment affects offspring physiology but that they will likely guide future investigations into the biological function of oocyte ETC and glycogen remodeling across species (particularly in flies, frogs, and humans where oocyte ETC and glycogen remodeling has already been described).

**Figure 6:**
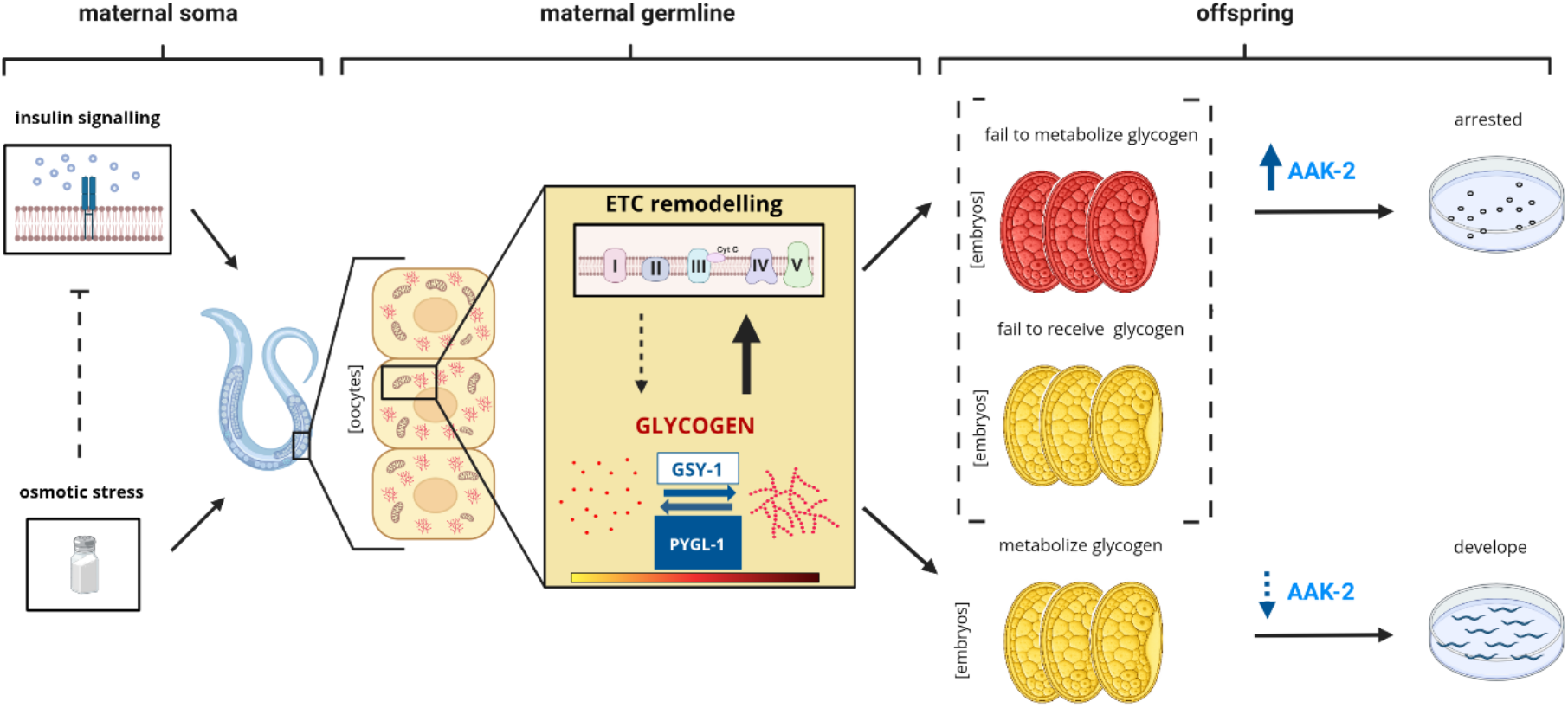
Proposed model for intergenerational adaptation to osmotic stress in *C. elegans*. Maternal exposure to mild osmotic stress and reduced maternal insulin/IGF-1 signaling both promote offspring development during periods of high osmotic stress by converging on the regulation of oocyte glycogen metabolism, which is also regulated by oocyte ETC function. Embryos that fail to metabolize (reduced *pygl-1* levels in maternal germline/early embryo) or receive (*isp-1* mutant animals) sufficient levels of glycogen have increased AAK-2 phosphorylation at Thr243 and fail to adapt to high levels of osmotic stress. Collectively, our findings indicate that the evolutionarily conserved phenomena of oocyte ETC and glycogen remodeling is responsive to a mother’s environment and that changes in oocyte ETC function and glycogen metabolism links multiple maternal inputs (maternal environment, maternal insulin signaling) to offspring’s future metabolism and response to stressors experienced by their mothers.

Like our findings in *C. elegans*, the original discovery of ETC/glycogen remodeling in *D. melanogaster* oocytes found that its onset is regulated by maternal insulin signaling^17^. In a follow up study in *D. melanogaster*, Hocaoglu *et al.*^56^, discovered that reduced insulin signaling in oocytes extended offspring survival under poor-nutrient conditions by 2.5-fold when compared to the offspring of control flies. This led to increased glycogen abundance in oocytes, similar to our findings here in *C. elegans.* However, they proposed a different mechanism of action for the observed effects on survival during nutrient restriction: that reduced ETC function in oocytes alters the oocyte NAD^+^/NADH ratio, which in turn alters the methionine cycle, histone methylation and gene expression in offspring. We cannot rule out that additional NAD^+^/NADH dependent pathways also function downstream of oocyte ETC remodeling in *C. elegans* (beyond our observed effects on glycogen) and that these additional pathways are also required to promote offspring’s future resistance to osmotic stress. Indeed, such additional pathways might explain why RNAi of *gsy-1* in germ cells does not completely eliminate adaptation in either the maternal exposure to osmotic stress or *daf-2* mutant models despite causing a substantial reduction in glycogen abundance in oocytes. Future studies will be critical in determining if additional metabolic changes in oocytes function with changes in glycogen abundance to mediate the link between a mother’s environment and offspring’s future physiology.

Despite these mechanistic differences across the *C. elegans* and *D. melanogaster* models, the collective findings raise the exciting possibility that oocyte ETC/glycogen remodeling is a “node” in oocytes that can regulate offspring’s response to multiple future stressors (osmotic stress, nutrient deprivation) and across multiple distantly related species (*C. elegans, D. melanogaster*) by altering the activity of specific downstream pathways in offspring. If true, then oocyte ETC/glycogen remodeling might represent more than a simple provisioning of resources, as previously speculated, but rather represent a mechanism that can regulate the activity of multiple downstream signaling pathways in early embryos and drive lasting changes in offspring physiology. This model is also consistent with the first observation of an oocyte-transmitted maternal effect in any animal, *B. mori* wintering diapause, which has similarly been mechanistically linked to oocyte sugar metabolism^14^. Future studies across paradigms will likely be critical in determining if different intergenerational stress paradigms might lead to stress specific patterns of ETC/glycogen remodeling in oocytes, or if different patterns of oocyte ETC/glycogen remodeling might influence multiple different downstream “receiver” signaling pathways in offspring. If so, such stress specific effects might explain why *C. elegans* intergenerational adaptation to osmotic stress prevents animals from simultaneously adapting to *P. vranovensis* infection^57^ or anoxia as observed in previous studies^33,58^ – *i.e*. this “node” in oocytes might be unable to promote adaptations to multiple stresses at the same time and “programming” offspring to adapt to one condition might come at the expense of their ability to respond to other stressors.

Beyond environmental stress paradigms, maternal age has also been proposed to alter offspring body size across species (humans, *C. elegans*, and *D. melanogaster*) via a mechanism that correlates with changes in oocyte mitochondrial morphology and that requires phosphorylation of AAK-2 in offspring^59^. While this effect does not involve a maternal environmental stress (and oocyte glycogen levels were not investigated) the mechanistic similarities raise the exciting possibility that the effects of maternal age on offspring development across species might also converge on this intergenerational signaling pathway. Together, these parallels across stress, nutrition and aging paradigms, as well as across invertebrate and vertebrate species, suggest that oocyte mitochondrial and glycogen remodeling may represent a broadly conserved phenomenon linking diverse maternal physiological states to offspring metabolic health, including in mammals, where parental metabolic exposures are already linked to offspring disease risk.

Several other classes of molecular mechanisms have been proposed to mediate multigenerational responses to stress in *C. elegans* including (but not limited to) small RNAs^60–62^, histone modifications^63–65^, N6-methyldeoxyadenine^64^, metabolite/yolk provisioning^57,66^, and changes in mtDNA DNA copy number^26^. Of these proposed mechanisms, changes in mtDNA copy number is the only other mechanism that implicates oocyte mitochondria. However, our findings indicate that mtDNA copy does not change in oocytes or embryos in our paradigm and that genes that regulate mtDNA copy number inheritance (*egl-20, mig-14*) were dispensable for this adaptation (**Fig 1**). Thus, the intergenerational signaling cascade we describe here (maternal insulin signaling, oocyte ETC/glycogen remodeling, and offspring AMP-kinase activity) appears to be a fundamentally novel mechanism linking a mother’s environment to adaptive changes in offspring physiology. Future investigations into whether the intergenerational signaling cascade we describe here intersects with other proposed mechanisms mediating intergenerational or transgenerational effects will likely be critical to broadening our understanding of how maternal environments can influence offspring’s future physiology.

Our proposed mechanism (**Fig 6**) centers on the abundance of oocyte glycogen stores and its ability to be metabolized in response to osmotic stress, which in turn affects the activity of AMP-kinase in offspring. Glycogen and AMP-kinase activity are linked through several mechanisms whose relative contributions remain to be fully resolved^45,46^. First, the beta subunit of AMP-kinase contains a glycogen binding domain, and glycogen has been proposed to directly bind and regulate AMP-kinase via allosteric effects^47^. Second, the glycogen binding domain of AMP-kinase regulates AMP-kinase localization and sequestration, thus it can affect the local substrates AMP-kinase has access to^47,67,68^. Lastly, glycogen can be metabolized to generate ATP, in turn regulating AMP-kinase via the indirect effects on the ATP/AMP ratio. In our paradigm there is a further possible mechanism of regulation: oocyte glycogen can be metabolized into osmolytes which would protect embryos against water loss induced by osmotic stress^3,69–71^. This could indirectly lower AMP-kinase activation in response to a given environmental osmolarity by lowering the effects of the external osmotic stress on cells. Our data favor a model in which the metabolism of oocyte-derived glycogen regulates offspring AMP-kinase phosphorylation (See Model in **Fig 6**) – by either increasing ATP production, increasing osmolyte production, or both. Specifically, maternal exposure to osmotic stress drives glycogen breakdown in oocytes (**Fig 3**) and this breakdown of glycogen (via PYGL-1) is required for offspring to bypass developmental arrest in response to future stress (**Fig 4**) and regulate the phosphorylation of AAK-2 (**Fig 5**). In addition, we note that *daf-2* mutants accumulate excess glycogen in oocytes and that the breakdown of this excess glycogen in embryos, again via PYGL-1, is sufficient for offspring to bypass developmental arrest when they are exposed to osmotic stress.

Collectively, these findings suggest that the ability to breakdown oocyte-derived glycogen (which can begin in oocytes in response to osmotic stress or in embryos if they receive excess glycogen from oocytes and are subsequently exposed to osmotic stress) is the mechanism that drives offspring resistance to future stress and that it does so in part by regulating offspring AMP-kinase activation. Moving forward, studies focused on offspring will be critical to determine the precise sequence of events in embryos by which glycogen breakdown causes changes in AMP-kinase phosphorylation. Similarly, future studies will also be critical in determining how changes AAK-2 activity in embryos intersects with ribosomal regulation (the ribosomal collision sensor GCN-1, eIF-2alpha phosphorylation, and ribosomal protein phosphorylation) to alter ribosomal function in arrested animals.

## Methods

### Strains

*C. elegans* strains were cultured and maintained at 20 °C unless noted otherwise. The Bristol strain N2 was utilized as the wild-type strain. Mutations used in this study:

|  | **Genotype** | **Source** |
|---|---|---|
| 1 | *N2 bristol* | CGC |
| 2 | *nuo-6*(qm200) | CGC |
| 3 | *nudf-7*(et19) | CGC |
| 4 | *gas-1*(fc21) | CGC |
| 5 | *mev-1*(kn1) | CGC |
| 6 | *isp-1*(qm150) | CGC |
| 7 | *egl-20*(n585) | CGC |
| 8 | *mig-14(*ga62) | CGC |
| 9 | *ymel-1*(tm1920) | CGC |
| 10 | *daf-2*(e1370) | CGC |
| 11 | *him-8*(e1489) | CGC |
| 12 | *eif-2α(S46A, S49A)* | CGC |
| 13 | *gcn-1(nc40)* | CGC |
| 14 | *gcn-1(n4827)* | CGC |
| 15 | *gcn-2(ok871)* | CGC |
| 16 | *gcn-2(ok886)* | CGC |
| 17 | *daf-16(m26) I* | CGC |
| 18 | *aak-2* (n6237I) | This study |
| 19 | *aak-2* (n6251) | This study |
| 20 | *aak-2 (n6252)* | This study |
| 21 | *aak-2 (bur1)* | This study |
| 22 | *aak-2 (ok524)* | This study |
| 23 | *aak-2 jbm54[C201S]* | This study |
| 24 | *aak-2 stj17[T243A])* | This study |
| 25 | *aak-2 (n6218)* | This study |
| 26 | *daf-16(m26) I; isp-1(qm150) IV.* | This study |

The nT1[qIs51] balancer was used to maintain *isp-1(qm150)* as a heterozygote in *daf-2*(e1370); *isp-1*(qm150/+).

Germline expression of *isp-1* was achieved utilizing sybIs7373 [P*pie-1*::*isp-1*::*tbb-2* 3’UTR]

Intestinal expression of *isp-1* was achieved utilizing P*ges-1*::*isp-1*::*tbb-2* 3’UTR

Neuronal expression of *isp-1* was achieved utilizing P*rgef-1*::*isp-1*::*tbb-2* 3’UTR and crossed to *isp-1* (qm150) for a genetic rescue.

For germline specific targeted RNAi studies the strains were used:

|  |
|---|
| *mkcSi13 [Psun-1::rde-1::tbb-2 3'UTR + unc-119(+)] II]; rde-1(mkc36) V* |
| *wago-3(pk1673); mkcSi13 [Psun-1::rde-1::tbb-2 3'UTR + unc-119(+)] II]; rde-1(mkc36) V* |
| *daf-2*(e1370); mkcSi13 [P*sun-1*::*rde-1*::tbb-2 3'UTR + *unc-119*(+) II].; *rde-1*(mkc36) V |
| *aak-2(T243A)*; *mkcSi13 [Psun-1::rde-1::tbb-2 3'UTR + unc-119(+) II].; rde-1(mkc36) V* |

For experimental crosses otIs [P*dat-1*::mCherry + P*ttx-3*::mCherry] was used to confirm cross progeny in experimental crosses as reported previously. ^3^

### isp-1 cloning

Full length ISP-1 cDNA (831 bp) along with its 5’ UTR (15 bp) and stop codons was PCR amplified using RNA isolated from WT *C. elegans* (Kuwahara et al., 2006). Primers included attB1 and attB2 sites, allowing for ISP-1 cDNA + 5’UTR to undergo BP recombination into pDONR221 [2,1]; this construct was sequence verified. Neuronal (rgef-1P: 3501 bp) and intestinal (ges-1p:2000 bp) promoters were amplified form WT genomic DNA. Primers included attB4 and attB1r sites, allowing for rgef-1P or ges-1 P to undergo BP recombination into pDONR P4-P1R [2,1]. Tbb-2 3 ‘UTR (323 bp) was amplified form WT genomic DNA. Primers included attB2r and attB3 sites, allowing for tbb-2 3’ UTR to undergo BP recombination into pDONR P2R-P3 [2,1]. Multi-site Gateway recombination was subsequently used to generate Pges-1::ISP-1::tbb-2 3 UTR and Prgef-1::ISP-1::tbb-2 3 UTR in the MosSCI pDEST backbone (pDEST R4-R3). All the constructs of BP and the final LR reactions were sequence verified.

### Injection methods

Gateway produced *isp-1* construct DNA was purified using Qiagen miniprep kit lacking RNAase in Buffer P1 (resuspension buffer) and eluted with nuclease free H2O. pPD93.73 (*myo-3*::GFP from ’95 Fire Vector Kit, expressed in body wall muscle) was used as an injection marker; pPD93.73 was linearized with PvuI and purified with Zymo cleanup kit. The injection mix contained 2ng/ul promoter::*isp-1* in pDest, linearized pPD93.73 at 5 ng/ul, and Bioruptor sheared Salmon testes DNA at 93 ng/ul for a total of 100 ng/ul in nuclease free H2O. DNA mix was microinjected into *isp-1* (qm150) mutants and injected animals were plated onto 60mm NGM cultures and incubated at 20 °C. After >1week, plates were screened for GFP+ F1s and these were cloned to new plates (all GFP+ F1s pooled per plate). Plates with GFP+ F2s were further assessed high penetrance and low mosaicism. Multiple lines per construct were tested for expression of *isp-1*::*tbb-2* UTR by semi-quantitative PCR and rescue of *isp-1* phenotypes.

### Osmotic stress assay

Adult animals raised under standard NGM conditions were collected in M9 buffer solution and embryos were isolated utilizing our embryo isolation protocol (see above). Embryos were then transferred to osmotically stressful cultures at various 834 concentrations (300, 350, and 400mM NaCl, etc) and incubated at room temperature for 48 hours. Hatched animals were then quantified and removed from the plate utilizing a standard laboratory vacuum suction system. Arrested animals were then collected using M9 buffer and allowed to recover in normal NGM culture conditions for 24 hours and then quantified. Osmotic stress assays using 500mM NaCl culture conditions were incubated for only 24 hours before quantification and recovery.

### Embryo isolation

Animals grown under standard NGM conditions were collected in M9 buffer and then washed 3x in diH20. Washed animals were then treated with a solution of 5% sodium hypochlorite and 5M NaOH, for 5 minutes under agitation. The embryos were then collected via centrifugation and treated with a secondary solution of 10% sodium hypochlorite for less than 1 minute and observed at 4x magnification to ensure successful embryo purification. Purified embryos were then collected and washed in 15 mL M9 and distributed to experimental cultures if the eggs were to be utilized in stress assays or washed in 15 mL diH2O, frozen in liquid nitrogen and stored at -80° C if embryos were to be utilized in proteomic or metabolomics analysis. *gas-1* mutants are extremely sensitive to sodium hypochlorite, needing up to 4 washes in 15 mL M9 (Supp figure 2b).

### Adaptation assay

Adolescent mothers (at the L4 larval stage) were collected in M9 buffer solution from standard NGM cultures, transferred to osmotically stressful 300mM NaCl cultures until animals were gravid (approximately 24 hours). After which time the cultures were collected and eggs were isolated using our egg isolation protocol. Harvested eggs were then transferred to 500mM NaCl cultures. After 48 hours hatched animals were quantified and removed utilizing vacuum suction and arrested embryos were collected in M9 buffer solution and allowed to recover in standard NGM culture conditions for 24 hours after which time recovered animals which regain normal activity were quantified.

### Experimental crosses

To generate heterozygous mutant strains genetic crosses were performed. Adolescent mothers (at the L4 larval stage) from the experimental or control strains were mated with males from the desired strains on NGM culture agar plates for 24-48 hours. The parent carrying the mutations were changed from maternal to paternal depending on the experimental design. Note that mitochondrial mutant males do not mate efficiently, and mitochondrial mutant mothers do not generate large brood sizes. To ensure successful mating males expressing fluorescent proteins (GFP or RFP) were used to confirm that progeny were actually produced from a successful mating. Laid eggs were then individually selected and transferred to 500mM NaCl cultures and allowed to develop at room temperature for 48 hours and the percentage of hatched animals was quantified.

### Oocyte dissections

Animals were raised to adolescence 795 (L4 stage) under standard culture conditions on NGM and then split into control or experimental cultures until day 1 of adulthood. Animals were then collected in M9 buffer and washed 3x. Single animals were then transferred to a microscope slide and dissected in a 10uL drop of paralytic 0.5 mg/mL tetramisole in M9. The procedure was performed using a dissection microscope at 4x magnification. The first incision was performed using a number 11 surgical scalpel on the paralyzed animal just posterior to the pharynx, exposing the intestine and the anterior gonad arm. The exposed gonad arm was then removed utilizing a 25 G x 5/8 needle and transferred to a 10uL solution of pure M9. The -1, -2, and -3 oocytes were then separated from the gonad arm using a 34 gauge RN needle, resulting the separation of the oocytes connected together with a small amount of gonadal sheath. These cells were then transferred via eyelash pick into 8uL of SMART seq solution for RNA sequencing, cold diH_2_O for proteomics, or cold 20% ethanol in diH_2_O for metabolomics and frozen at -80 °C.

### Low input oocyte protein digests for proteomics

384-well plates (Part No. 0030129547, Eppendorf, Hamburg, Germany) containing oocytes were thawed to 4 °C over 30 min and then centrifuged at 600 ×g for 20 s. The Tecan UNO (Mannedorf, Switzerland) single-cell dispenser was then used to dispense 1 μL of solution containing 5 ng of trypsin and 5 ng of Lys-C from the Promega (Madison, Wisconsin) Rapid-Digestion Trypsin/Lys-C Kit (Part No. VA1061) into each well. The proteases were dissolved in the Rapid Digest Buffer included in the kit. The following settings were used for the Tecan UNO: fluid group was set to aqueous, and fluid class was set to “aqueous (surfactant free). The well plates were then covered with a silicone sealing mat (ArcticWhite), again centrifuged at 600 ×g for 20 s, wrapped in aluminum foil and incubated in a humidified chamber for 1 h at 70 °C. The plate was then cooled to 4 °C for 15 min and again centrifuged as described above. The silicone sealing mat was replaced with adhesive storage foil (Eppendorf, H 819 amburg, Germany) and samples were then stored at -20 °C. C-18 column (Affinisep AttractSPE tips -Le Houlme, France) clean-up and desalting for low volume samples (less than 15ug) was then performed to remove contaminants that could interfere with mass spectrometry instruments. Briefly, frozen samples were reconstituted in 0.1% TFP in LCMS grade H2O and sonicated in a water bath for 5 minutes. The samples were then spun on a C-18 column and centrifuged 1,000 x g for 2 minutes. The column was then washed with 0.1% formic acid in LCMS grade H2O. The sample was then eluted from the column in 0.1% formic acid in 70% ACN in LCMS H2O and centrifuged 1,000 x g for 2 minutes. The samples were then dried in a SpeedVac at 30° C for 2 hours. Dried samples were then resuspended in 0.1% formic acid in LCMS grade H2O at a concentration of 2ug/uL for LC-MS/MS analysis.

### Embryo Proteomics

*C.elegans* samples were homogenized on the Bead Ruptor Elite (Cat#19-042E, Omni International) for 30 s in 4% SDS solution containing 1x HALT Protease made available under a CC-BY-NC-ND 4.0 (Cat# 78442, Thermo Fisher Scientific). Samples 886 were sonicated and clarified via centrifugation and transferred to a Protein LoBind Eppendorf tube. Proteins were quantified using the Pierce BCA Protein Assay Kit (Cat# 23227, Thermo Fisher Scientific) and 100 μg of protein was aliquoted for digestion. Protein digestion utilized the S-Trap (Cat# CO2-Mini, Protifi) platform to remove any SDS prior to LCMS/MS analysis. Briefly proteins were reduced with Dithiothreitol (DTT) for 20 minutes, alkylated with Iodoacetamide (IAA) for 20 minutes and digested overnight with Trypsin/Lys-C (Cat# V5072, Promega) at a ratio of 50:1 (protein:enzyme (w/w)). Peptides were eluted off the S-Trap Column and dried down in a Genevac SpeedVac. Dried samples were then cleaned up with C-18 reverse phase columns from Harvard Apparatus(Cat# 74-4601) and dried down before resuspension for LC-MS/MS analysis. Dried samples were resuspended in 50 μL 0.1% FA (LS118-1, Fisher Scientific) and diluted with 50 μL of 0.1% TFA (LS119-500, Fisher Scientific). Data Independent acquisition analyses were performed on Orbitrap Eclipse coupled to Vanquish Neo system (Thermo Fisher Scientific). The FAIMS Pro source (Thermo Fisher 900 Scientific) was located between the nanoESI source and the mass spectrometer. 2 μg of digested peptides were separated on a nano capillary column (20 cm × 75 μm I.D., 365 μm O.D., 1.7 μm C18, CoAnn Technologies, Washington, # HEB07502001718IWF) at 300 nL/min. Mobile phase A consisted of LC/MS grade H2O (LS118-500, Fisher Scientific), mobile phase B consisted of 20% LC/MS grade and H2O and 80% LC/MS grade acetonitrile (LS122500, Fisher Scientific), and both mobile phases contained 0.1% formic acid. The LC gradient was: 1% B to 24% B in 110 min, 85% B in 5 min, and 98% B for 5 min, with a total gradient length of 120 min. For FAIMS, the selected compensation voltage (CV) was applied (−40V, -55V, -70V) throughout the LC-MS/MS runs. Full MS spectra were collected at 120,000 resolution (full width half-maximum; FWHM), and MS2 spectra at 30,000 resolutions (FMWH). Both the standard automatic 910 gain control (AGC) target and the automatic maximum injection time were selected. A precursor range of 380-980 m/z was set for MS2 scans, and an isolation window of 50 m/z was chosen with a 1 m/z overlap for each scan cycle. 32% HCD collision energy was used for MS2 fragmentation. To generate a hybrid library for directDIA™ analysis in Spectronaut, pooled samples underwent data dependent acquisition employing 11 distinct FAIMS CV settings ranging from -30 to 80 CV. Full MS spectra were collected at 120,000 resolution (full width half-maximum; FWHM), and MS2 spectra at 30,000 resolutions (FMWH). Both the standard automatic gain control (AGC) target and the automatic maximum injection time were selected. Ions were filtered with charge 2–5. An isolation window of 1.6m/z was used with quadruple isolation mode. Ions were fragmented using higher-energy collisional dissociation (HCD) with a collision energy of 30%. DIA data was processed in Spectronaut (version 18, Biognosys, Switzerland) using direct DIA. Data was searched against a *Caenorhabditis elegans* reference proteome including both UniProt and TrEMBL databases. The manufacturer’s default parameters were used. Briefly, trypsin/P was set as digestion enzyme and two missed cleavages were allowed. Cysteine carbamidomethylation was set as fixed modification, and methionine oxidation and protein N-terminus acetylation as variable modifications. Identification was performed using a 1% q value cutoff on precursor and protein levels. Both peptide precursors and protein false discovery rate (FDR) were controlled at 1%. Ion chromatograms of fragment ions were used for quantification. For each targeted ion, the area under the curve between the XIC peak boundaries was calculated. To enhance proteome coverage, the DDA raw files were utilized in Library Extension Runs to generate a hybrid library.

### Phosphoproteomics

Proteins were extracted from *C. elegans* samples at room temperature for 30 seconds on the Bead Ruptor Elite (Omni International) in a 4% SDS solution containing 1X Halt Protease and Phosphatase Inhibitor. Protein concentration was then measured using the Pierce BCA Protein Assay. Subsequently, the proteins underwent tryptic digestion and purification using C18 columns. For global proteome quantitation, 10 % of peptide mixture were then resuspended in 0.1% TFA at a concentration of 1µg/ul to be injected on the instrument. For global phosphoproteome quantitation. 90% of peptide mixture was carried out a phosphopeptide enrichment using Magnetic Ti-IMAC beads according to manufacturer protocols. Briefly, the peptide mixture was reconstituted in 200 µl loading buffer 0.1 M glycolic acid in 80% CAN and 5% TFA and vortex. The mixture was transferred to MagReSyn® Ti-IMAC HP microparticles (Resyn bioscience) and incubated for 20 min at room temperature with slow mixing. The supernatant was removed using magnetic separator. The beads were sequentially washed using 2x with Buffer 1 (80% ACN, 1% TFA) and 1x with Buffer 2 (10% ACN, 0.2% TFA). The bound phosphopeptides were eluted with 150 µl elution buffer (1% NH4OH) for 10 min. The eluates were dried down and further cleaned up using C18 columns for the LC-MS/MS. For an unbiased quantitative proteome and phosphoproteome, data-independent acquisition (DIA) approach was employed uisng on Orbitrap Eclipse coupled to Vanquish Neo system (Thermo Fisher Scientific). The FAIMS Pro source (Thermo Fisher Scientific) was located between the nanoESI source and the mass spectrometer. 2 μg of digested peptides were separated on a nano capillary column (20 cm × 75 μm I.D., 365 μm O.D., 1.7 μm C18) at 300 nL/min. The LC gradient was: 1% B to 26% B in 51 min, 80% B in 5 min, and 98% B for 4 min, with a total gradient length of 60 min. Full MS spectra were collected at 120,000 resolution (full width half-maximum; FWHM), and MS2 spectra at 30,000 resolutions (FMWH). Both the standard automatic gain control (AGC) target and the automatic maximum injection time were selected. A precursor range of 380-980 m/z was set for MS2 scans, and an isolation window of 50 m/z was chosen with a 1 m/z overlap for each scan cycle. 32% HCD collision energy was used for MS2 fragmentation. DIA data was processed in Spectronaut (version 18) using a direct DIA mode. Data was searched against C.elegans reference proteomes The manufacturer’s default parameters were used. Identification was performed using a 1% q-value cutoff on precursor and protein levels. Both peptide precursors and protein false discovery rate (FDR) were controlled at 1%.

### LIMMA Proteomics analysis

Differential abundance of proteins was performed using LIMMA and eBayes tools. For differential abundance of proteomics based on 8 replicates of samples (Fig. 1D), proteins with 0 variance and proteins with >30% missingness in any experimental group were removed from analysis. We examined the remaining missingness in the dataset and found that proteins that had missing values in any samples had lower abundance values than proteins without any missingness. Since the missing values were not missing at random, we used a left-censored imputation method (impute.QRILC). Data were then log2 transformed to prepare for LIMMA analysis. We next used LIMMA eBayes to analyze the log2 transformed data with custom contrasts. Both LIMMA and empirical Bayes were performed using robust methods to avoid one protein or sample having too much influence over the results. The results were multiple testing corrected using the Benjamini-Hochberg method to maintain a 5% false discovery rate. For proteomics comparisons based on 3 biological replicates (Fig. 1B, 1C, 5A), only proteins that exhibited zero variance across all samples were also removed from the analysis. The remaining data were then transformed via variance normalization stabilization to prepare for the LIMMA eBayes analysis. We next used LIMMA eBayes to analyze the normalized data with custom contrasts. Both LIMMA and empirical Bayes were performed using robust methods to avoid one protein or sample having too much influence over the results. Proteins were scored as significant if the exhibited a >2-fold change in abundance between conditions and *p* < 0.05. We note that previous studies of oocyte proteomics have normalized to either Complex IV, which we and others have found to not change in abundance in response to stress, or to total mitochondrial protein^17,18^. Consistent with this, protein abundance in oocyte proteomics replicates was normalized to Complex IV (COX-6A and COX-6B) abundance to control for technical variation between samples. This method was validated in **Fig. 1B** accurately detect changes in NDUF-7 abundance in *nduf-*958 *7* mutant animals. In addition, for samples containing >20 oocytes, we found that the proteins detected included some proteins known to be highly expressed in gonadal sheath cells but not expressed in oocytes such as TBH-1, TDC-1, INX-8 and IFY-1 (**Supplemental Table 2**). By contrast, many other gonadal sheath specific proteins were undetectable such as LIM-6, LIM-7, FKH-6, LIN-11, APX-1, MOM-2, and SYS-1 (**Supplemental Table 2**). These findings are consistent with the fact that some gonadal sheath is unable to be detached from 3-oocyte samples by dissection. However, that the amount of gonadal sheath present in our samples is likely small and thus only highly abundant proteins present in sheath tissue were detectable – we estimate under 10% of total biomass from our dissections comes from attached sheath tissue – see **Fig. 1A**. Nonetheless, it is possible that variation in the amount of attached gonadal sheath might serve as a source of variation between low-input samples dissected from different animals. Thus, to control for any differences in the relative abundance of gonadal sheath between samples we normalized all samples to the abundance of ASB-2, which we and others have found to be highly abundant in mitochondria derived from somatic cells including gonadal sheath cells but absent from mitochondria derived from germ cells such as oocytes (**see Supplemental Fig. 1A**).

### Embryo Metabolomics

Animals for metabolomics were grown as previously described unless otherwise noted. Starved animals represent animals placed on NGM agar plates under standard laboratory conditions. For metabolomics analysis, an acetonitrile-methanol-water (4:4:2 v/v) extraction was used. Samples were homogenized in ice-cold 4:4:2 acetonitrile (A955, Fisher):methanol (A456, Fisher):water (W6, Fisher) such that final extraction solvent to sample ratios were 2 million cells per 1 mL of extraction solvent. After solvent addition, extracts were vortexed for 10 s, sonicated in a water bath for 5 min, and incubated on wet ice for 1 hour. Incubation was followed by a 10-minute centrifugation at 17,000 g and 4°C to collect the solid protein pellet at the bottom of the tube and separate it from the metabolite-containing supernatant. Next, 1.6 million cell equivalents were aliquoted from the supernatant, dried under vacuum, and resuspended in 200 uL LCMS-grade water. 50ul of this resuspension was taken for GC-MS analysis. For GCMS analysis samples were subjected to derivatization with 30 μL of methoxyamine (11.4 mg/mL) in pyridine and 70 μL of MTBSFA+1%TMCS TBDMS for GCMS analysis on an Agilent 7890GC/5977bMSD as described previously^72,73^. The oven program was: initial 95°C ramp to 118°C at 40°C/min then hold for 2 min, then ramp to 300°C at 12°C/min, and hold at 300°C for 5 min. A neat standard of glycerol was derivatized and used to determine retention time (10.77min) and fragment ions (170.8, 188.7, 289, and 377). 170.8m/z was used as the quantifier ion, and others used to qualify the peak. Injection volume was 1 uL. Data were analyzed in Skyline.

### Low input oocyte RNA-seq and analysis

RNA was prepped from oocytes using the Direct-zol RNA microprep kit (ZYMO Research) and was sequenced using an Illumina NovaSeq 6000. Reads were aligned to the WBcel235 reference genome and quantified with STARsolo. A gene matrix composed of a row for each gene and column for each sample was built and brought into R using SummarizedExperiment. Genes with fewer than ten reads across all samples were removed. Differential analysis of the quantified genes was performed with DESeq251.

### Droplet digital PCR

The mtDNA copy number was measured with droplet digital PCR (ddPCR). Animals were lysed as described above and diluted two-fold and 2 uL of diluted lysate was combined with 0.25 µL of 10-µM of each oligonucleotide primer. Mixtures of lysate and primer were combined with 12.5 µL of Bio-Rad QX200TM ddPCR EvaGreen Supermix and nuclease-free water to a volume of 25 µL in EppendorfTM 96-well twin.tecTM PCR plates. The samples were loaded on the Bio-Rad Droplet Generator and processed using the standard droplet generation program for EvaGreen samples. PCR was performed with an annealing temperature of 58°C. The samples were loaded on the Bio-Rad QX200 Droplet Reader and mtDNA-carrying droplets was quantified using the QX manager software. The following primers were utilized to measure mtDNA. Primer MP435:5′-GTCCTTGTGGAATGGTTGAATTTAC-3′ Primer MP436:5′-GTACTTAATCACGCTACAGCAGC-3′ Gitschlag et al., 2020

### RNAi

RNAi was administered through feeding animals with RNAi-expressing HT115 *E. coli* using standard methods and libraries^74,75^. Knockdown utilizing RNAi was achieved by feeding animals transformed HT115 that express a double stranded RNA (dsRNA) that targets the genes of interest. NGM agar solution with tetracycline was used to select for the L440 RNAi plasmid in the HT115 bacteria and IPTG (Isopropyl β-D-1-thiogalactopyranoside) was added to strongly activate T7 polymerase expression to enhance transcription of dsRNA. RNAi plates were seeded with the transformed bacterial culture, dried and allowed to grow overnight. *C. elegans* mothers were collected and embryos were extracted, isolated and sterilized as described above. Animals were scored using our lugol’s staining protocol and imaged at day 1 of adulthood.

### Lugol’s stain

Animals were collected in M9 and washed 3x by gravity sedimentation. Staining and imaging quickly is critical because lugol’s iodine stain fades quickly in *C. elegans* tissue. A 1:20 dilution of Logul’s iodine in M9 was added to washed animals and was incubated for 5 minutes. Animals were washed 3x in M9 by centrifugation (13,000 rpm for 10 seconds) and the washed pellet was transferred to glass slides to be imaged by bright field at 60x oil immersion using an Olympus BX51 microscope equipped with an Amscope MU1403 camera. Approximately 20 images were collected per condition.

### Lugol’s stain quantification

To quantify the changes in both color and intensity within oocytes and embryos, we developed an analysis pipeline combining QuPath (v0.6.0) and Python (v3.14). Individual cells were manually annotated in QuPath. For oocyte images, annotations were also made to indicate the developmental stage order. The resulting annotations were then exported as labeled PNG masks using a custom Groovy script. Cellular color and intensity were analyzed using custom Python scripts. The original RGB images were converted to the HSV (hue, saturation, value) color space using scikit-image. For each annotated cell, we calculated the mean hue, saturation, and value. To account for phase wraparound in circular hue angles, the mean hue was calculated using the scipy.stats.circmean function. Additionally, the intensity of each cell was calculated using the equation:

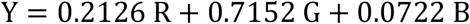

where R, G, and B, represent the intensities in the red, green, and blue channels, respectively. To measure color shifts relative to the control group, we calculated the hue difference ΔH from the control (H_control_) using the equation:

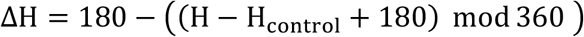

The modulo operation accounts for the angular boundary wrapping between 360° and 0°. Additionally, we inverted the sign of the phase shift so that positive shifts denote a red shift while negative shifts denote a yellow shift. To mitigate batch effects across experimental days, the hue shift was calculated relative to the baseline mean hue of the controls acquired on the same day.

#### Code Availability

All code and instructions related to this project can be found at: https://github.com/vaioic/burton-lab-lugols-analysis.

### Western blotting

Animals grown on respective RNAi plates were collected at day-1 of adulthood and embryos were isolated by hypochlorite treatment and plated onto standard NGM (50 mM NaCl) plates or on high salt (400 mM) plates. After 6 hours of exposure, embryos were collected in M9, centrifuged and resuspended in Buffer C (50 mM HEPES, 100 mM Kcl, 1mM MgCl2, 1mM EGTA, 10 percent glycerol) solution with protease inhibitors (Sigma P2714) and phosphatase inhibitors (Sigma 4906845001) and lysed by sonication using a Diagenode Bioruptor. Protein levels were then quantified using a standard BCA assay and samples were diluted to ensure equal levels of protein were loaded. Laemmli SDS Sample Buffer with beta-mercaptoethanol were added to each sample. Samples were incubated at 95 °C for 10 minutes and then frozen until loading. Samples were boiled, loaded, and run in a 12% gel for SDS-PAGE. The gel was transferred onto a nitrocellulose membrane for 1.5 hours followed by blocking with 5% BSA in 0.1% TBST 1 hour at room temperature. Membranes were then probed with primary antibodies (diluted in blocking buffer) overnight at 4 C. Phospho-AMPK-1 was detected using the rabbit anti-phospho-AMPK-1 antibody (Cell Signaling Technology 2535S; 1:5000 dilution) and actin was detected using a mouse anti-actin antibody (Sigma A5441; 1:5000 dilution). The blots were washed 3 times in 0.1% TBST for 3 minutes each followed by secondary antibody incubation for 2 hours at RT. Following secondary antibodies were diluted in 0.1% TBST and used: anti-rabbit secondary antibody (Invitrogen SA5-35571; 1:10,000 dilution) and anti-mouse secondary antibody (Invitrogen A-21057; 1:10,000 dilution).

### Fluorescence microscopy

Epifluorescence images of *asb-2*(syb9897)::RFP were obtained using a Leica M250FA stereoscope with a Hamamatsu ORCA-ER camera and LAS-X1024 software using a RFP filter set. To image expression in oocytes, an adult day 2 animal was dissected in M9 (see dissection protocol above) and oocytes from this animal were transferred to a 4uL drop of M9 on a cytology microscope slide and a coverslip was immediately applied. This side was then quickly imaged using the Leica M250FA microscope (less than 2 minutes) at 63x magnification. Fluorescence intensity was quantified by measuring average intensity across the oocytes using FIJI software.

### Semi-Quantitative PCR

For each well, 2 worms were pooled in 4 ul standard lysis buffer containing 1 mg/mL proteinase K (2-4 wells were run per genotype per PCR). Samples were flash frozen and lysed at 65 deg for 10 min followed by 85 deg for 1 minute to inactivate ProK. mRNA transcripts were produced using Maxima H Minus Reverse Transcriptase kit (Thermo Scientific), with lysates briefly treated with dsDNase enzyme prior to reverse transcription (2 ul lysate into 10ul final reaction volume) following standard protocols and including both Oligod(T) and random hexamer primers. cDNAs were diluted 1:1 with H2O and 2 ul was used as template in 30 ul PCR reactions (DreamTaq, Thermo Scientific). Reactions were divided and run at 25, 30, and 35 cycles and compared to *cdc-42* (housekeeping gene) as well as positive and negative controls using the following primers:

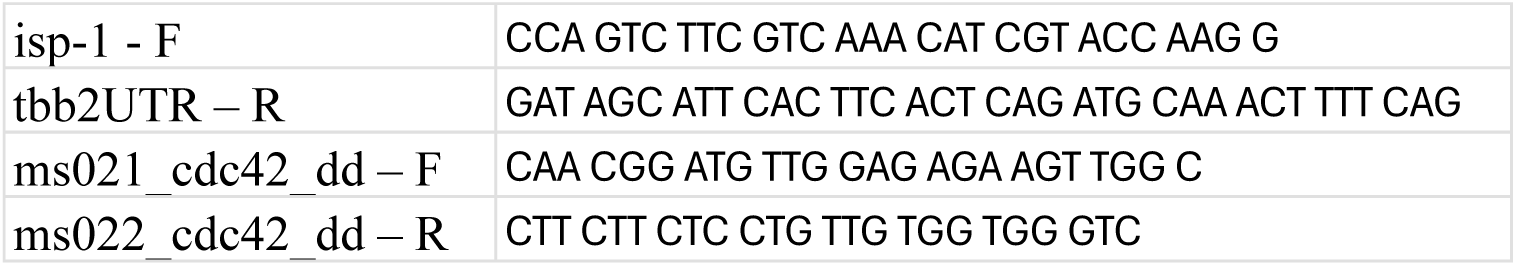

### Statistical analysis

One-way ANOVA analysis was used for Figs. 2C, 2D, 2E, 2F, 3C, 3E, 3G, 4A, 4D, 4E, 4F, 4H, 5C, and Supplementary Figs 3A, 3B, 3C, 3F, 4A, 4B, 4D, 4E.

Two-way ANOVA analysis with Šídák’s or Tukey’s multiple comparison test was used for Figs. 1I, 2B, 2G, 2H, 3H, 4I, 4J, 5B, 5E, 5F, 5G, 5I, 5J and Supplementary Figs 3D, 3H, 3I, 5C, 5D.

Unpaired two-tailed Student’s t-test with multiple hypothesis correction was used for Figs. 1H and Supplementary Fig. 1D.

For Fig. 2I, J and Fig. 5 D, data was analyzed using logistic regression with Firth’s bias correction in R version 4.6.1 (CRAN). To test whether the proportion of worms that developed differed by genotype, we fit a logistic regression model with Firth’s bias correction, which accommodates genotypes for which no worms developed (1). Pairwise genotype comparisons were performed using emmeans, with Tukey adjustment for multiple comparisons (2).

1. Kosmidis I (2026). *brglm2: Bias Reduction in Generalized Linear Models*. R package version 1.1.1, https://CRAN.R-project.org/package=brglm2.
2. Lenth R, Piaskowski J (2026). *emmeans: Estimated Marginal Means, aka Least-Squares Means*. R package version 2.0.5, https://rvlenth.github.io/emmeans/.

LIMMA analysis for global proteomics (Figs. 1 and 4) and DESeq2 analysis for RNA-seq (Supplementary Fig. 1) are described above in separate sections.

No statistical method was used to predetermine sample size. The experiments were not randomized. The investigators were not blinded to allocation during experiments and outcome assessment

## Data availability

RNA-seq data that support the findings of this study have been deposited at NCBI GEO and are available under the accession code GSE261225. Proteomics data that support the findings of this study have been deposited in the PRIDE repository and are available under the accession code PXD050192 and PXD055906.

## Acknowledgments

We thank D. Chandler for helpful comments on the manuscript text and the *Caenorhabditis* Genetics Stock Center (funded by the NIH National Center for Research Resources) and National BioResource Project (NRBP) for strains. We also thank Noah Lancaster and Josh Coon for help guiding proteomics method development, and H. Robert Horvitz who provided assistance in the identification of *aak-2* in initial screens. We also thank Noah Lancaster and Joshua Coon of the National Center for Quantitative Biology of Complex Systems (NIH GM108538) for helpful discussions and support. This work was supported by funds from the VAI MeNu research grant program. A.P.W. and C.O. were supported by NIH R01NS092558. We also thank diverse members of the VAI core facilities including M. Adams and B. Fillinger for assistance with single oocyte RNA-seq and analysis. Model image was created with BioRender.com.

## Author Contributions

J.F.C, U.R and N.O.B conceived the project. J.F.C, U.R. K.N., D.G. D.W., C.C., C.O., M.S., A.G., K.N., X.A., and D.W. performed the experiments. C.O. and A.W. designed and generated *aak-2* mutant strains. E.W., X.A., R.K., and H.L. oversaw and analyzed the proteomics and J.W.T., U.R. and J.F.C analyzed the Lugol’s staining. R.K., M.P., R.S., A.W., and N.O.B. oversaw individual experiments and analyzed the data. N.O.B. oversaw the overall project and wrote the manuscript.

---

**Supp. Figure 1:**
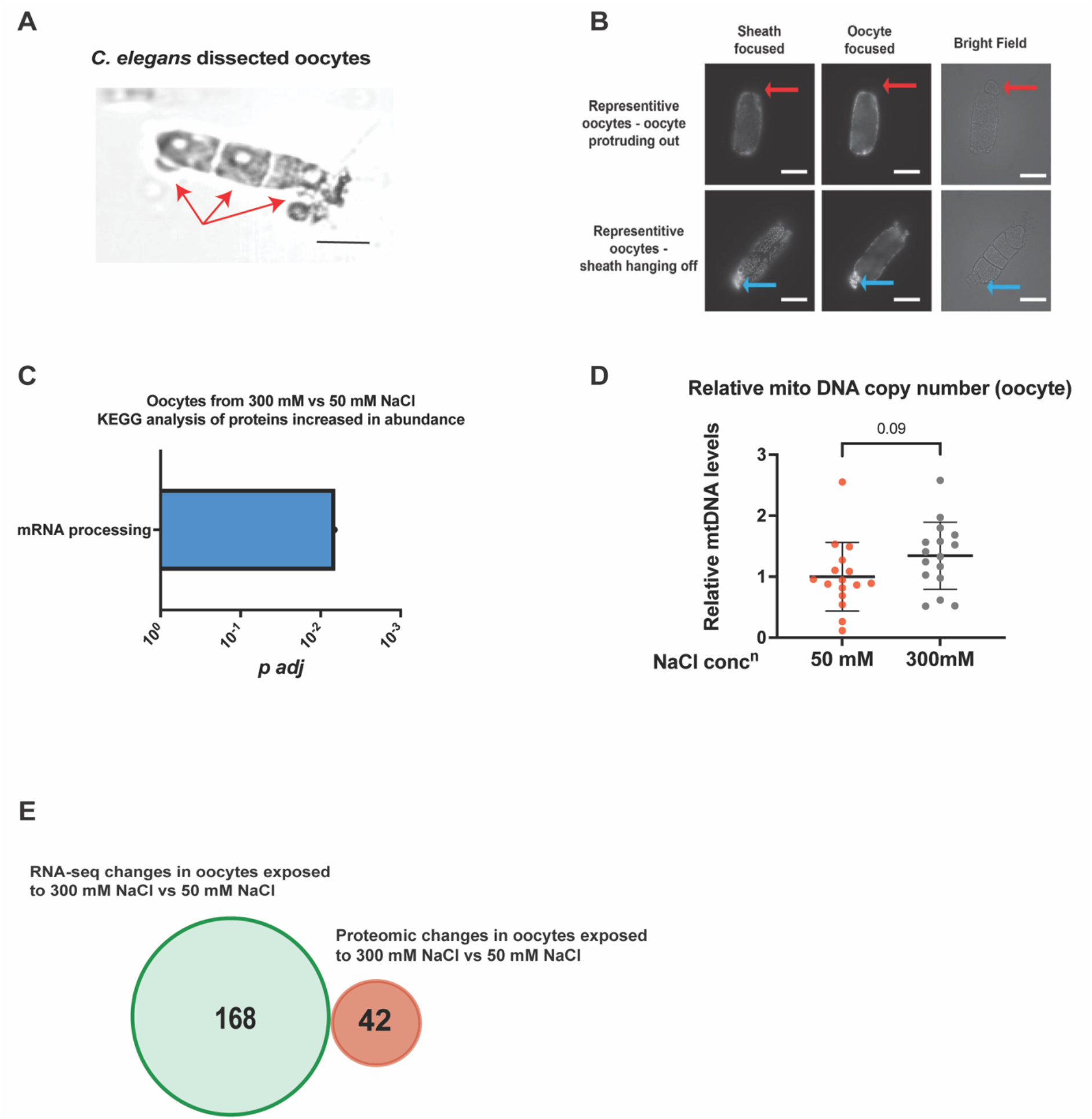
Changes in maternal oocyte ETC protein abundance in response to osmotic stress is independent of changes in oocyte mRNA, interreference from sheath cell or oocyte mtDNA copy number. **A.** Representative image of -1, -2, and -3 oocytes dissected from an individual animal. Scale bar 30 μM. **B.** Representative images of an endogenously tagged *ASB-2::mCherry* in oocytes and associated sheath cells. Red arrows point to an oocyte that has bulged out of the surrounding sheath cells and you can see that there is zero fluorescence emitted. Blue arrows point to sheath cells attached to dissected oocytes which show high *ASB-2::mCherry* expression. These findings are consistent with RNA expression studies finding that asb-2 is expressed exclusively in somatic cells and not in germ cells. The bulging out oocyte also highlights that the background fluorescence observed in the oocyte focused samples comes from bleed through from surrounding sheath cells. Scale bars 50 μm. **C.** Biological compartment analysis of proteins altered in abundance in oocytes from parents exposed to 300 mM NaCl compared to wild-type oocytes from parents exposed to 50 mM NaCl using WormEnrichr. [**chen** et al, **Kuleshov** et al]. **D.** Relative mitochondrial DNA copy number was determined using digital-droplet PCR (ddPCR) in dissected oocytes from *C. elegans* mothers exposed to 50 mM NaCl or to 300 mM NaCl (mild osmotic stress) for 24 hours. n -16 replicates (12 oocytes per replicate). Unpaired students’ *t*-test with Welch correction. error bars -s.d. **E.** Venn diagram of mRNAs and proteins that changed in abundance in wild-type oocytes from animals exposed to 50 mM NaCl versus 300 mM NaCl for 24 hours. Source data are provided in Supplemental table 8.

**Supp. Figure 2:**
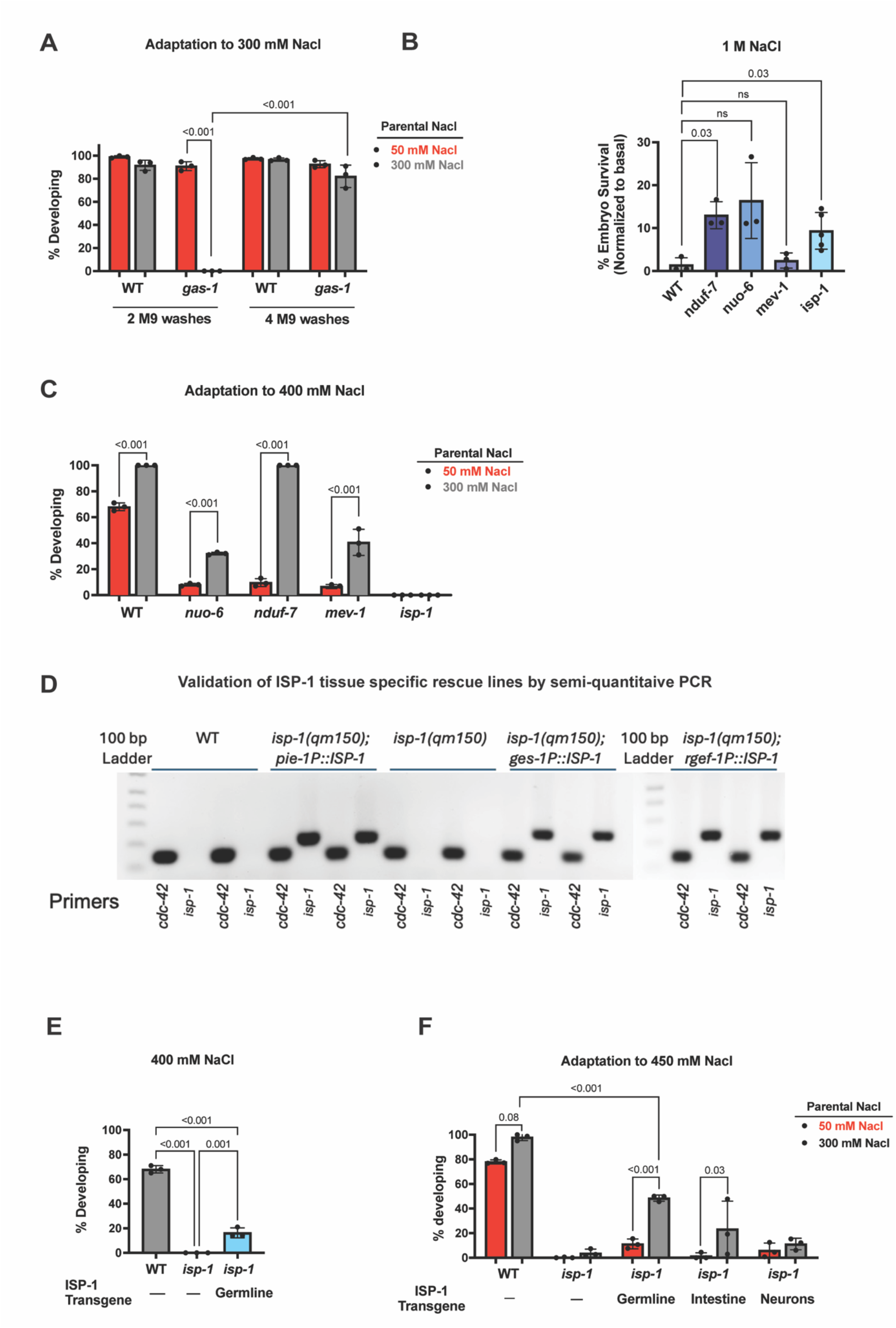
The electron transport chain (ETC), particularly complex III, function regulates animals’ response to osmotic stress. **A.** Percent of WT and *gas-1(fc21)* embryos that were mobile and developing after 48 hours of exposure to 300 mM NaCl after maternal exposure to 50 mM or 300mM NaCl. The embryos were washed (after egg prep with sodium hypochlorite) either 2 or 4 times with M9 before exposing them to 300 mM NaCl. n = 3 replicates of > 100 offspring per replicate, Two-way ANOVA with Tukey’s multiple comparison test, error bars – s.d. **B.** Percent of WT, *nduf-7(et19)*, *nuo-6(qm200)*, *mev-1(kn1)* and *isp-1(qm150)* embryos surviving after 48 hours on 1 M NaCl. The embryo survival at 1 M NaCl is normalized to embryo survival at standard 50 mM NaCl conditions for each strain respectively n = 3 replicates of > 75 offspring per replicate, one-way ANOVA, error bars – s.d. **C.** Percent of WT, *nduf-7(et19)*, *nuo-6(qm200)*, *mev-1(kn1)* and *isp-1(qm150)* embryos that were mobile and developing after 48 hours of exposure to 400 mM NaCl after maternal exposure to 50 mM or 300mM NaCl. n = 3 replicates of > 40 offspring per replicate, Two-way ANOVA with Šídák’s multiple comparison test, error bars – s.d. **D.** Validation of transgenic ISP-1 expression by quantitative Real time PCR (qRT-PCR) in WT, *isp-1(qm150), isp-1(qm150); pie-1p::isp-1*(Germline ISP-1 rescue)*, isp-1(qm150); ges-1p::isp-1* (Intestinal ISP-1 rescue), and *isp-1(qm150); rgef-1p::isp-1* (Neuronal ISP-1 rescue). *cdc-42* was used as a control. **E.** Percent of WT, *isp-1(qm150), isp-1(qm150); pie-1p::isp-1*(Germline ISP-1 rescue) embryos that were mobile and developing after 48 hours of exposure to 400 mM NaCl. n = 3 replicates of > 50 offspring per replicate, one-way ANOVA, error bars – s.d. **F.** Percent of WT, *isp-1(qm150), isp-1(qm150); pie-1p::isp-1*(Germline ISP-1 rescue)*, isp-1(qm150); ges-1p::isp-1* (Intestinal ISP-1 rescue), and *isp-1(qm150); rgef-1p::isp-1* (Neuronal ISP-1 rescue) embryos developing after 48 hours of exposure to 450 mM NaCl after maternal exposure to 50 mM or 300mM NaCl. n = 3 replicates of > 40 offspring per replicate, Two-way ANOVA with Tukey’s multiple comparison, error bars – s.d. vSource data are provided in Supplemental table 8.

**Supp. Figure 3:**
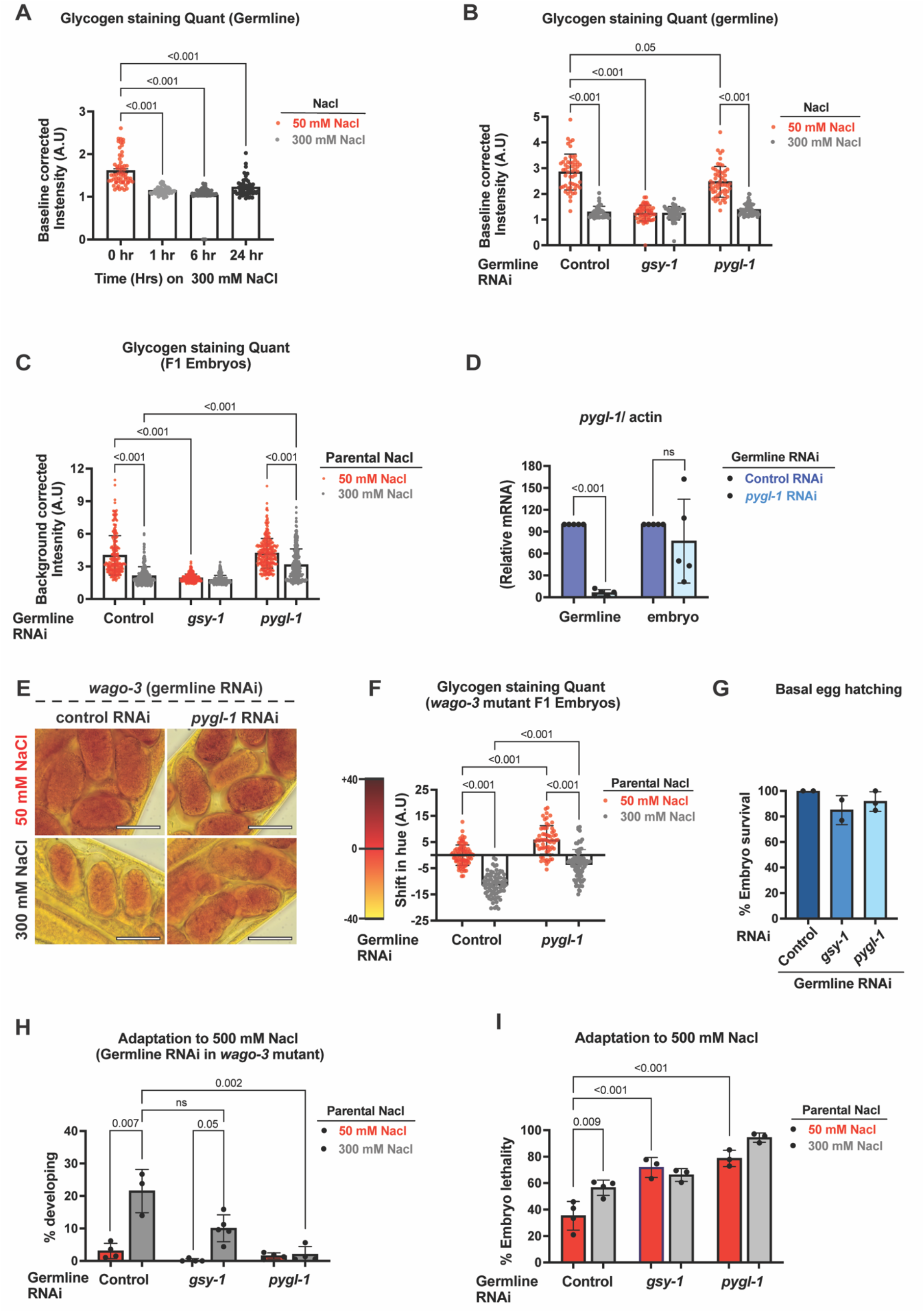
Maternal oocyte glycogen responds to osmotic stress and oocyte glycogen metabolism is required for intergenerational adaptation to osmotic stress. **A.** Quantification of background corrected intensity for Fig. 3A. n=3 replicates, 20 animals imaged per replicate, one-way ANOVA, error bars – s.d. **B.** Quantification of background corrected intensity for Fig. 3D. n=3 replicates, 20 animals imaged per replicate, one-way ANOVA, error bars – s.d. **C.** Quantification of background corrected intensity for Fig. 3F. n=3 replicates, 20 animals imaged per replicate, one-way ANOVA, error bars – s.d. **D.** Relative *pygl-1* mRNA levels were determined by digital droplet PCR (ddPCR). *sun-1p::rde-1; rde-1(mkc36)* (Germline specific-RNAi) animals were grown on control or *pygl-1* RNAi till day-1 of adulthood. The germlines were dissected and in parallel F1 embryos were collected, lysed and prepped for making cDNA. ddPCR was performed for both germline and embryo samples*. pygl-1* mRNA levels were normalized to as *actin*. **E.** Representative brightfield images of lugol’s iodine-stained embryos from *wago-3(pk1673)*; *sun-1p::rde-1; rde-1*(*mkc36*) (Germline specific-RNAi) worms. Worms were grown on control or *pygl-1* RNAi at standard conditions (50 mM NaCl) and either exposed to 50 mM or mild salt stress (300 mM Nacl) for 24 hours followed by lugol’s staining of respective F1 embryos in uterus of these mothers. **F.** Quantification of supp. Fig. 3E. Shift in hue is calculated with respect to worms grown on control RNAi at 50 mM NaCl. n = 3 replicates, >15 animals imaged per replicate, one-way ANOVA, error bars – s.d. **G.** Percent of basal embryos hatching. *sun-1p::rde-1; rde-1*(*mkc36*) worms were grown on control, *gsy-1* or *pygl-1* RNAi at standard conditions (50 mM NaCl) and embryos from these mothers were scored and allowed to develop at standard conditions. The animals that developed were scored and percent of embryo hatching was calculated. N = 3 replicates of > 100 offspring per replicate, One-way ANOVA. error bars -s.d. **H.** Percent of embryos developing at 500 mM NaCl. *wago-3(pk1673)*; *sun-1p::rde-1; rde-1*(*mkc36*) worms were grown on control, *gsy-1* or *pygl-1* RNAi at standard conditions (50 mM NaCl) and at early day-1 of adulthood were either exposed to 50 mM or mild salt stress (300 mM Nacl) for 24 hours. The embryos from these mothers were exposed to 500 mM NaCl for 48 hours, following which developing and mobile larva were scored. n = 3 or 4 replicates of > 80 offspring per replicate, Two-way ANOVA with Šídák’s multiple comparison test. error bars -s.d. **I.** Percent of embryo lethality at 500 mM NaCl for Fig 3H, where *sun-1p::rde-1; rde-1*(*mkc36*) (Germline specific-RNAi) worms were grown on control, *gsy-1* or *pygl-1* RNAi at standard conditions (50 mM NaCl) and at early day-1 of adulthood were either exposed to 50 mM or mild salt stress (300 mM Nacl) for 24 hours. The embryos from these mothers were exposed to 500 mM NaCl for 48 hours, following which developing and mobile larva were scored and the arrested eggs were transferred and scored. n = 3 or 4 replicates of > 80 offspring per replicate, Two-way ANOVA with Tukey’s multiple comparison test. error bars -s.d. Scale bars: 30 μM Source data are provided in Supplemental table 8.

**Supp. Figure 4:**
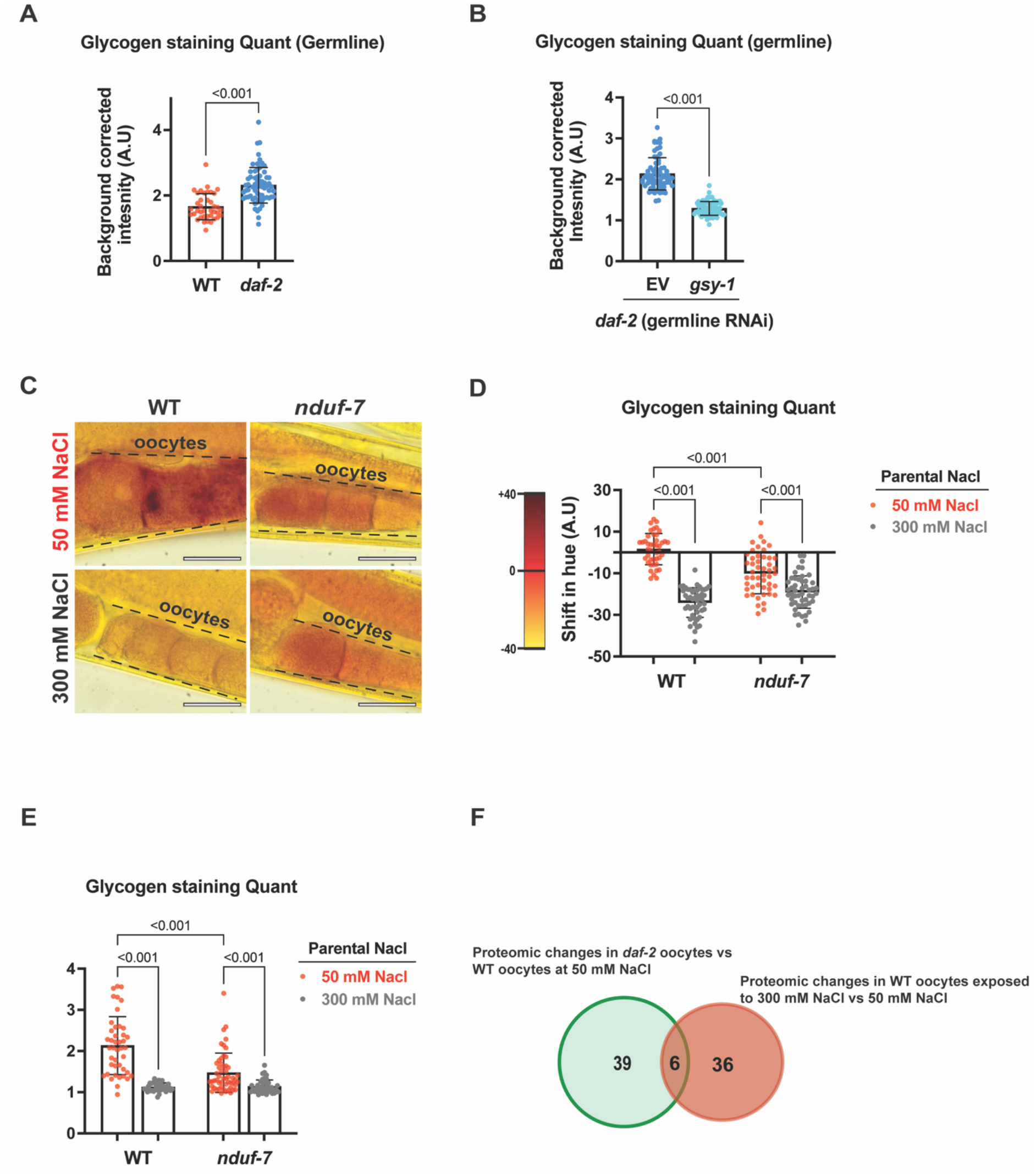
Reduced insulin signaling increase oocyte glycogen levels while disruption in ETC function reduce oocyte glycogen levels. **A.** Quantification of background corrected intensity for Fig. 4A. n=3 replicates, 20 animals imaged per replicate, Unpaired students’ *t*-test with Welch correction, error bars – s.d. **B.** Quantification of background corrected intensity for Fig. 4C. n=3 replicates, 20 animals imaged per replicate, Unpaired students’ *t*-test with Welch correction, error bars – s.d. **C.** Representative brightfield images of lugol’s iodine-stained oocytes of WT and *nduf-7(et19)* adult worms. Worms were grown at standard conditions (50 mM NaCl) and at early day-1 of adulthood either exposed to 50 mM or mild salt stress (300 mM Nacl) for 24 hours followed by lugol’s staining. **D.** Quantification of Supp. Fig. 4C. Shift in hue is calculated for each condition with respect to WT at standard condition (50 mM NaCl). n=3 replicates, >12 animals imaged per replicate, one-way ANOVA, error bars – s.d. **E.** Quantification of background corrected intensity for Fig. 4C. n=3 replicates, >12 animals imaged per replicate, one-way ANOVA, error bars – s.d. **F.** Venn diagram of proteins that changed in abundance in oocytes of WT versus *daf-2(e1370)* day-1 adult animals. Experiments were carried out at 20° C. Scale bars: 30 μM Source data are provided in Supplemental table 8.

**Supplemental Figure 5:**
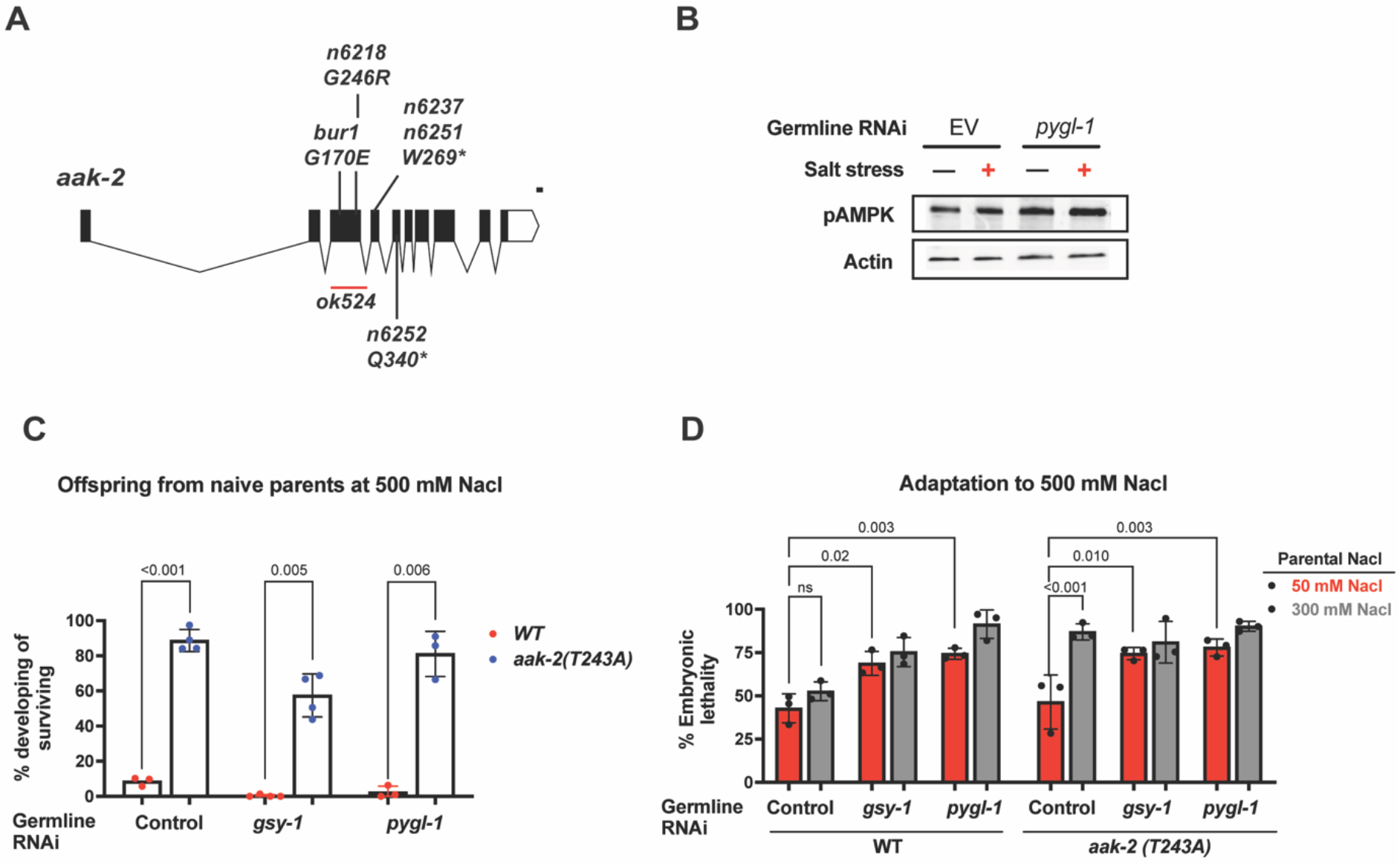
Maternal glycogen metabolism modulates offspring AMPK activity, while *aak-2* mutants bypass osmotic stress-induced developmental arrest. **A.** Schematic of the *aak-2* gene depicting different *aak-2* mutations recovered from mutagenesis screens. Red bar represents deleted region. **B.** Western blot for analysing relative pAMPK levels. *sun-1p::rde-1; rde-1(mkc36)* (Germline specific-RNAi) animals were grown on control or *pygl-1* RNAi till day-1 of adulthood. The embryos were isolated and exposed to 50 mM or 400 mM NaCl for 6 hours. Animals were then collected and embryos extracted for western blot analysis. Actin was probed as a loading control. The quantification is as shown in Fig. 5E. **C.** Percent of embryos developing at 500 mM NaCl. *sun-1p::rde-1; rde-1*(*mkc36*) (Germline specific-RNAi) and *aak-2(T243A); sun-1p::rde-1; rde-1*(*mkc36*) worms were grown on control, *gsy-1* or *pygl-1* RNAi at standard conditions (50 mM NaCl) and embryos from these mothers were exposed to 500 mM NaCl for 48 hours, following which developing and mobile larva was scored and % developing worms were plotted with respect to total number of survivors. n = 4 replicates of > 80 offspring per replicate, Two-way ANOVA with Šídák’s multiple comparison test. error bars -s.d **D.** Percent of embryos developing at 500 mM NaCl. *sun-1p::rde-1; rde-1*(*mkc36*) (Germline specific-RNAi) and *aak-2(T243A); sun-1p::rde-1; rde-1*(*mkc36*) worms were grown on control, *gsy-1* or *pygl-1* RNAi, at standard conditions (50 mM NaCl) and at early day-1 of adulthood were either exposed to 50 mM or mild salt stress (300 mM Nacl) for 24 hours. The embryos from these mothers were exposed to 500 mM NaCl for 48 hours, following which developing and mobile larva was scored and % developing worms were plotted with respect to total number of eggs. n = 4 replicates of > 80 offspring per replicate, Two-way ANOVA with Šídák’s multiple comparison test. error bars -s.d

## References

1. Weismann, A., Parker, W. N. (William N. & Rönnfeldt, H. The Germ-Plasm; a Theory of Heredity. (New York, Scribner’s, 1893).

2. Luna, E., Bruce, T. J. A., Roberts, M. R., Flors, V. & Ton, J. Next-Generation Systemic Acquired Resistance. Plant Physiol. 158, 844–853 (2012).

3. Burton, N. O. et al. Insulin-like signalling to the maternal germline controls progeny response to osmotic stress. Nat. Cell Biol. 19, 252–257 (2017).

4. Rechavi, O. et al. Starvation-Induced Transgenerational Inheritance of Small RNAs in C. elegans. Cell 158, 277–287 (2014).

5. Willis, A. R. et al. A parental transcriptional response to microsporidia infection induces inherited immunity in offspring. Sci. Adv. 7, eabf3114 (2021).

6. Dantzer, B. et al. Density Triggers Maternal Hormones That Increase Adaptive Offspring Growth in a Wild Mammal. Science 340, 1215–1217 (2013).

7. Huypens, P. et al. Epigenetic germline inheritance of diet-induced obesity and insulin resistance. Nat. Genet. 48, 497–499 (2016).

8. Painter, R. C., Roseboom, T. J. & Bleker, O. P. Prenatal exposure to the Dutch famine and disease in later life: An overview. Reprod. Toxicol. 20, 345–352 (2005).

9. Cheng, Q. et al. Prenatal and early-life exposure to the Great Chinese Famine increased the risk of tuberculosis in adulthood across two generations. Proc. Natl. Acad. Sci. 117, 27549–27555 (2020).

10. Hales, C. N. & Barker, D. J. P. Type 2 (non-insulin-dependent) diabetes mellitus: the thrifty phenotype hypothesis. Diabetologia 35, 595–601 (1992).

11. Liu, Y. et al. Caloric restriction prevents inheritance of polycystic ovary syndrome through oocyte-mediated DNA methylation reprogramming. Cell Metab. 37, 920–935.e6 (2025).

12. Aiken, C. E. & Ozanne, S. E. Transgenerational developmental programming. Hum. Reprod. Update 20, 63–75 (2014).

13. Simmons, R. A., Templeton, L. J. & Gertz, S. J. Intrauterine Growth Retardation Leads to the Development of Type 2 Diabetes in the Rat. Diabetes 50, 2279–2286 (2001).

14. Yamashita, O., Yaginuma, T. & Hasegawa, K. Hormonal and Metabolic Control of Egg Diapause of the Silkworm, Bombyx mori (Lepidoptera: Bombycidae). Entomol. Gen. 195–211 (1981) doi:10.1127/entom.gen/7/1981/195.

15. Burton, N. O. & Greer, E. L. Multigenerational Epigenetic Inheritance: Transmitting information across generations. Semin. Cell Dev. Biol. 127, 121–132 (2022).

16. Fitz-James, M. H. & Cavalli, G. Molecular mechanisms of transgenerational epigenetic inheritance. Nat. Rev. Genet. 23, 325–341 (2022).

17. Sieber, M. H., Thomsen, M. B. & Spradling, A. C. Electron Transport Chain Remodeling by GSK3 during Oogenesis Connects Nutrient State to Reproduction. Cell 164, 420–432 (2016).

18. Rodríguez-Nuevo, A. et al. Oocytes maintain ROS-free mitochondrial metabolism by suppressing complex I. Nature 607, 756–761 (2022).

19. Wang, C. et al. High-throughput label-free single-cell proteomics enabled by multicolumn NanoLC. Nat. Commun. 17, 11066 (2026).

20. Rauthan, M., Ranji, P., Abukar, R. & Pilon, M. A Mutation in Caenorhabditis elegans NDUF-7 Activates the Mitochondrial Stress Response and Prolongs Lifespan via ROS and CED-4. G3 GenesGenomesGenetics 5, 1639–1648 (2015).

21. Formosa, L. E., Dibley, M. G., Stroud, D. A. & Ryan, M. T. Building a complex complex: Assembly of mitochondrial respiratory chain complex I. Semin. Cell Dev. Biol. 76, 154–162 (2018).

22. Ugalde, C., Janssen, R. J. R. J., van den Heuvel, L. P., Smeitink, J. A. M. & Nijtmans, L. G. J. Differences in assembly or stability of complex I and other mitochondrial OXPHOS complexes in inherited complex I deficiency. Hum. Mol. Genet. 13, 659–667 (2004).

23. Urso, S. J., Comly, M., Hanover, J. A. & Lamitina, T. The O-GlcNAc transferase OGT is a conserved and essential regulator of the cellular and organismal response to hypertonic stress. PLoS Genet. 16, e1008821 (2020).

24. Hibshman, J. D. & Goldstein, B. LEA motifs promote desiccation tolerance in vivo. BMC Biol. 19, 263 (2021).

25. Harding, H. P. et al. An integrated stress response regulates amino acid metabolism and resistance to oxidative stress. Mol. Cell 11, 619–633 (2003).

26. Zhang, Q. et al. The memory of neuronal mitochondrial stress is inherited transgenerationally via elevated mitochondrial DNA levels. Nat. Cell Biol. 23, 870–880 (2021).

27. Yang, W. & Hekimi, S. Two modes of mitochondrial dysfunction lead independently to lifespan extension in Caenorhabditis elegans. Aging Cell 9, 433–447 (2010).

28. Ishii, N. et al. A mutation in succinate dehydrogenase cytochrome b causes oxidative stress and ageing in nematodes. Nature 394, 694–697 (1998).

29. Feng, J., Bussière, F. & Hekimi, S. Mitochondrial electron transport is a key determinant of life span in Caenorhabditis elegans. Dev. Cell 1, 633–644 (2001).

30. Kayser, E. B., Morgan, P. G. & Sedensky, M. M. GAS-1: a mitochondrial protein controls sensitivity to volatile anesthetics in the nematode Caenorhabditis elegans. Anesthesiology 90, 545–554 (1999).

31. Anglas, U. et al. MAK-2 Kinase Is Required for Extended Longevity and Enhanced Stress Resistance Resulting From Mild Impairment of Mitochondrial Function in isp-1 Mutants. Aging Cell 25, e70537 (2026).

32. Bayer, E. A. et al. Insulin signaling and osmotic stress response regulate arousal and developmental progression of C. elegans at hatching. Genetics 220, iyab202 (2022).

33. Frazier, H. N. & Roth, M. B. Adaptive sugar provisioning controls survival of C. elegans embryos in adverse environments. Curr. Biol. CB 19, 859–863 (2009).

34. Cid, E., Geremia, R. A., Guinovart, J. J. & Ferrer, J. C. Glycogen synthase: towards a minimum catalytic unit? FEBS Lett. 528, 5–11 (2002).

35. Agius, L. Role of glycogen phosphorylase in liver glycogen metabolism. Mol. Aspects Med. 46, 34–45 (2015).

36. Singh, M., et al. Glycogen supports glycolytic plasticity in neurons. Proc. Natl. Acad. Sci. 122, e2509003122 (2025).

37. Zou, L. et al. Construction of a germline-specific RNAi tool in C. elegans. Sci. Rep. 9, 2354 (2019).

38. Schreier, J., Pshanichnaya, L., Kielisch, F. & Ketting, R. F. A genetic framework for RNAi inheritance in Caenorhabditis elegans. EMBO Rep. 26, 4072–4099 (2025).

39. Burton, N. O. et al. Neurohormonal signaling via a sulfotransferase antagonizes insulin-like signaling to regulate a Caenorhabditis elegans stress response. Nat. Commun. 9, 5152 (2018).

40. Carman, L., Schuck, R. J., Li, E. & Nelson, M. D. An AMPK biosensor for Caenorhabditis elegans. MicroPublication Biol. 2022, (2022).

41. Hawley, S. A. et al. Characterization of the AMP-activated protein kinase kinase from rat liver and identification of threonine 172 as the major site at which it phosphorylates AMP-activated protein kinase. J. Biol. Chem. 271, 27879–27887 (1996).

42. Lee, H. et al. The Caenorhabditis elegans AMP-activated Protein Kinase AAK-2 Is Phosphorylated by LKB1 and Is Required for Resistance to Oxidative Stress and for Normal Motility and Foraging Behavior. J. Biol. Chem. 283, 14988–14993 (2008).

43. Zmijewski, J. W. et al. Exposure to hydrogen peroxide induces oxidation and activation of AMP-activated protein kinase. J. Biol. Chem. 285, 33154–33164 (2010).

44. Carman, L., Schuck, R. J., Li, E. & Nelson, M. D. An AMPK biosensor for Caenorhabditis elegans. 10.17912/MICROPUB.BIOLOGY.000596 (2022)

45. Herzig, S. & Shaw, R. J. AMPK: guardian of metabolism and mitochondrial homeostasis. Nat. Rev. Mol. Cell Biol. 19, 121–135 (2018).

46. Garcia, D. & Shaw, R. J. AMPK: Mechanisms of Cellular Energy Sensing and Restoration of Metabolic Balance. Mol. Cell 66, 789–800 (2017).

47. McBride, A., Ghilagaber, S., Nikolaev, A. & Hardie, D. G. The Glycogen-Binding Domain on the AMPK β Subunit Allows the Kinase to Act as a Glycogen Sensor. Cell Metab. 9, 23–34 (2009).

48. Reduced glycogen availability is associated with increased AMPKα2 activity, nuclear AMPKα2 protein abundance, and GLUT4 mRNA expression in contracting human skeletal muscle. https://cdnsciencepub.com/doi/10.1139/h06-003 (2026).

49. Watt, M. J. et al. β-adrenergic stimulation of skeletal muscle HSL can be overridden by AMPK signaling. FASEB J. 18, 1445–1446 (2004).

50. Jobava, R. et al. Adaptive translational pausing is a hallmark of the cellular response to severe environmental stress. Mol. Cell 81, 4191–4208.e8 (2021).

51. Held, J. P. et al. A tRNA processing enzyme is a key regulator of the mitochondrial unfolded protein response. eLife 11, e71634 (2022).

52. Oltion, K. et al. An E3 ligase network engages GCN1 to promote the degradation of translation factors on stalled ribosomes. Cell 186, 346–362.e17 (2023).

53. Costa-Mattioli, M. & Walter, P. The integrated stress response: From mechanism to disease. Science 368, eaat5314 (2020).

54. Pakos-Zebrucka, K. et al. The integrated stress response. EMBO Rep. 17, 1374–1395 (2016).

55. Zheng, X. et al. Exploring the multifaceted role of GCN1: Implications in cellular responses and disease pathogenesis. Biomed. Pharmacother. Biomedecine Pharmacother. 175, 116681 (2024).

56. Hocaoglu, H., Wang, L., Yang, M., Yue, S. & Sieber, M. Heritable shifts in redox metabolites during mitochondrial quiescence reprogram progeny metabolism. Nat. Metab. 3, 1259–1274 (2021).

57. Burton, N. O. et al. Intergenerational adaptations to stress are evolutionarily conserved, stress-specific, and have deleterious trade-offs. eLife 10, e73425 (2021).

58. Dey, S., Proulx, S. R. & Teotónio, H. Adaptation to Temporally Fluctuating Environments by the Evolution of Maternal Effects. PLOS Biol. 14, e1002388 (2016).

59. Maternal aging increases offspring adult body size via transmission of donut-shaped mitochondria | Cell Research. https://www.nature.com/articles/s41422-023-00854-8.

60. Houri-Ze’evi, L., Teichman, G., Gingold, H. & Rechavi, O. Stress Resets Transgenerational Small RNA Inheritance. 669051 Preprint at 10.1101/669051 (2019).

61. Houri-Zeevi, L., Teichman, G., Gingold, H. & Rechavi, O. Stress resets ancestral heritable small RNA responses. eLife 10, e65797 (2021).

62. Ow, M. C. & Hall, S. E. Inheritance of Stress Responses via Small Non-Coding RNAs in Invertebrates and Mammals. Epigenomes 8, 1 (2023).

63. Kishimoto, S., Uno, M., Okabe, E., Nono, M. & Nishida, E. Environmental stresses induce transgenerationally inheritable survival advantages via germline-to-soma communication in Caenorhabditis elegans. Nat. Commun. 8, 14031 (2017).

64. Wan, Q.-L. et al. N6-methyldeoxyadenine and histone methylation mediate transgenerational survival advantages induced by hormetic heat stress. Sci. Adv. 7, eabc3026 (2021).

65. Fabrizio, P., Garvis, S. & Palladino, F. Histone Methylation and Memory of Environmental Stress. Cells 8, 339 (2019).

66. Aupérin, N. et al. Soma-to-germline miRNA inheritance through yolk promotes stress resilience in progeny. Nat. Struct. Mol. Biol. 33, 985–997 (2026).

67. Oligschlaeger, Y. et al. The recruitment of AMP-activated protein kinase to glycogen is regulated by autophosphorylation. J. Biol. Chem. 290, 11715–11728 (2015).

68. Janzen, N. R., Whitfield, J. & Hoffman, N. J. Interactive Roles for AMPK and Glycogen from Cellular Energy Sensing to Exercise Metabolism. Int. J. Mol. Sci. 19, 3344 (2018).

69. Possik, E. et al. FLCN and AMPK Confer Resistance to Hyperosmotic Stress via Remodeling of Glycogen Stores. PLOS Genet. 11, e1005520 (2015).

70. Possik, E. & Pause, A. Glycogen: A must have storage to survive stressful emergencies. Worm 5, e1156831 (2016).

71. Davis, M., Montalbano, A., Wood, M. P. & Schisa, J. A. Biphasic adaptation to osmotic stress in the C. elegans germ line. Am. J. Physiol. -Cell Physiol. 312, C741–C748 (2017).

72. Ma, E. H. et al. Metabolic Profiling Using Stable Isotope Tracing Reveals Distinct Patterns of Glucose Utilization by Physiologically Activated CD8+ T Cells. Immunity 51, 856–870.e5 (2019).

73. Kaymak, I. et al. Carbon source availability drives nutrient utilization in CD8+ T cells. Cell Metab. 34, 1298–1311.e6 (2022).

74. Kamath, R. S. et al. Systematic functional analysis of the Caenorhabditis elegans genome using RNAi. Nature 421, 231–237 (2003).

75. Rual, J.-F. et al. Toward improving Caenorhabditis elegans phenome mapping with an ORFeome-based RNAi library. Genome Res. 14, 2162–2168 (2004).

76. Kuleshov, M. V. et al. Enrichr: a comprehensive gene set enrichment analysis web server 2016 update. Nucleic Acids Res. 44, W90–97 (2016).

77. Chen, E. Y. et al. Enrichr: interactive and collaborative HTML5 gene list enrichment analysis tool. BMC Bioinformatics 14, 128 (2013).

